# Benchmarking AlphaFold for protein complex modeling reveals accuracy determinants

**DOI:** 10.1101/2021.10.23.465575

**Authors:** Rui Yin, Brandon Y. Feng, Amitabh Varshney, Brian G. Pierce

## Abstract

High resolution experimental structural determination of protein-protein interactions has led to valuable mechanistic insights, yet due to the massive number of interactions and experimental limitations there is a need for computational methods that can accurately model their structures. Here we explore the use of the recently developed deep learning method, AlphaFold, to predict structures of protein complexes from sequence. With a benchmark of 152 diverse heterodimeric protein complexes, multiple implementations and parameters of AlphaFold were tested for accuracy. Remarkably, many cases had highly accurate models generated as top-ranked predictions, greatly surpassing the performance of unbound protein-protein docking, whereas antibody-antigen docking was largely unsuccessful. While AlphaFold-generated accuracy predictions were able to discriminate near-native models, previously developed scoring protocols improved performance. Our study demonstrates that end-to-end deep learning can accurately model transient protein complexes, and identifies areas for improvement to guide future developments to reliably model any protein-protein interaction of interest.

## Introduction

Protein-protein interactions are the basis of many critical and fundamental cellular and molecular processes, including inhibition or activation of enzymes, cellular signaling, and recognition of antigens by the adaptive immune system. High resolution structural characterization of these interactions provides insights into their molecular basis, as well as structure-guided design of binding affinities and identification of inhibitors. However, structures for large numbers of molecular interactions remain undetermined experimentally, due to limitations in resources, and the challenges of structural determination techniques.

In response to this need, numerous predictive computational methods to model structures of protein-protein complexes have been developed over several decades, including protein docking methods that use unbound or modeled component structures as input to perform rigid-body global searches in six dimensions [1-5], and template-based modeling methods that generate models of complexes based on known structures [6, 7]. Challenges for docking algorithms include side chain and backbone conformational changes between unbound and bound structures, large search spaces, and inability to capture key energetic features in grid-based and other rapidly computable functions, leading to false positive models among top-ranked models or lack of any near-native models within large sets of predicted models. Developments such as explicit side chain flexibility during docking searches [8], use of normal mode analysis to represent protein flexibility [9, 10], clustering [11, 12] or re-scoring [13-16] docking models to improve ranking of near-native models, and use of experimental data as restraints for docking [17] have led to some improvement in docking success, and examples of these and other advances specifically designed to address the challenge posed by protein backbone flexibility are highlighted in a recent review [18]. However, the Critical Assessment of Predicted Interactions (CAPRI) blind docking prediction experiment [19] and several protein docking benchmarks [20, 21], which have enabled the systematic assessment of predictive docking performance, revealed persistent shortcomings of current computational docking approaches. Several protein-protein complex targets had no accurate model generated by any teams in a set of recent CAPRI rounds [22], while benchmarking of multiple docking algorithms in 2015 showed no accurate models within sets of top-ranked predictions for many of the test cases [20]. A more recent benchmarking study with 67 antibody-antigen docking test cases highlighted the limited success for current global docking approaches, which was more pronounced for cases with more conformational changes between unbound and bound structures [23].

The recently developed AlphaFold algorithm (AlphaFold v. 2.0) uses an end-to-end deep neural network to generate structural models from sequence [24], showing unprecedentedly high modeling accuracy and substantially surpassing the performance of other teams in the most recent Critical Assessment of Structural Prediction (CASP) round [25]. An important element of that algorithm is the use of multiple sequence alignment (MSA) based features to infer residue contacts, building on previous work demonstrating the use of co-evolution in contact prediction [26, 27]. Its reported capability to model homomultimers [24], as well as a recently reported adaptation of AlphaFold to enable modeling of heteroprotein assemblies [28], raises the question of how accurately AlphaFold can model transient heteroprotein complexes, including classes of complexes that have challenged previously developed and currently available docking approaches. As the AlphaFold deep learning model was trained using experimentally determined structures of individual protein chains [24], and its accuracy was partly enabled by residue contacts within tertiary structures inferred from multiple sequence alignment, it is not clear whether it can reliably generate protein-protein interface structures, due to their distinct physicochemical properties versus inter-residue interactions within proteins, as well as lack of explicit MSA signal from pairs of residues across the protein-protein interface in the sequences.

Here we report a systematic assessment of the accuracy of AlphaFold in performing end-to-end modeling of transient protein complexes, using 152 heterodimeric test cases from Protein-Protein Docking Benchmark version 5.5 (BM5.5) [20, 23] which represent three previously established docking difficulty levels, and classes of interactions including enzyme-containing complexes, antibody-antigen complexes, as well as a range of other complex types. Comparing performance of two implementations of AlphaFold with a global protein docking algorithm, ZDOCK [29] showed remarkable and superior accuracy across the benchmark, even with only five models generated per test case. Determinants of modeling success were assessed by case category and other features, and a number of scoring functions, in addition to pTM and pLDDT scores generated by AlphaFold, were tested to find optimal scoring criteria to identify correct docking models from AlphaFold. These results illustrate that while not successful for all cases and complex types, AlphaFold is a powerful tool for complex modeling, showing the power and advantage of end-to-end deep learning versus previous docking approaches. Our results also highlight areas for future optimization and developments in this framework, or other end-to-end deep learning frameworks, to effectively and reliably model most or all transient protein-protein complexes.

## Results

### Performance of AlphaFold on protein-protein complex prediction

To assess the accuracy of AlphaFold in predicting structures of transient protein-protein complexes, we used Protein-Protein Docking Benchmark 5.5 (BM5.5) [20, 23], which contains complexes spanning many classes of interactions that were identified from the Protein Data Bank [30] using an automated pipeline followed by manual inspection and curation. All heterodimeric protein-protein complexes from that benchmark were identified for this analysis, corresponding to 152 test cases (**Table S1**). Sequences of the two chains from each test case were input to AlphaFold, which generated structural models of the protein complexes using unpaired MSAs, without the use of templates. Additionally, the “advanced” interface in ColabFold [28] was utilized to generate protein complex models using the AlphaFold framework. ColabFold uses different databases and MSA generation algorithm, but its speed and web accessibility make it an useful alternative to a locally installed full AlphaFold pipeline. To permit comparison with a current docking approach, the rigid-body docking program ZDOCK (version 3.0.2) [29] and the IRAD scoring function [31] were used to perform global docking and rank models for all complexes, using unbound protein structures as input.

The performance of AlphaFold, ColabFold, and ZDOCK was assessed by comparison of models with experimentally determined structures of the bound complexes; overall success rate comparisons are shown in **Figure 1a**, for top 1 (T1) and top 5 (T5) ranked models for each test case, with per-case performance shown in **Figure S1**. Remarkably, AlphaFold was able to generate models with Acceptable or higher accuracy, based on CAPRI criteria [22], for approximately half of the test cases, and for most of those cases, Medium or High accuracy level models were generated. Additionally, the top-ranked model (T1) based on AlphaFold pTM score represented the highest accuracy level for each case, leading to only a modest improvement in success when allowing five predicted models per case (T5). The success rate for ColabFold was similar to the success of AlphaFold, indicating that the different sequence databases and MSA procedure did not reduce or otherwise alter the capability of the AlphaFold deep learning model to generate near-native complex models. Inspection of per-case performance (**Figure S1**) confirmed that ColabFold and AlphaFold success was highly correlated across the test cases. Rigid-body global docking success from ZDOCK was considerably lower than AlphaFold and ColabFold, particularly for Medium and High accuracy models, although a subset of cases were successful for ZDOCK while not successfully predicted by AlphaFold or ColabFold (**Figure S1**). A representative successfully modeled complex from AlphaFold is shown in comparison with the experimentally determined structure of the complex (PDB code 2×9A; E coli TolA/Phage G3B complex) in **Figure 1b**, demonstrating modeling of a virus-host protein-protein interaction with atomic-level accuracy.

**Figure 1.**
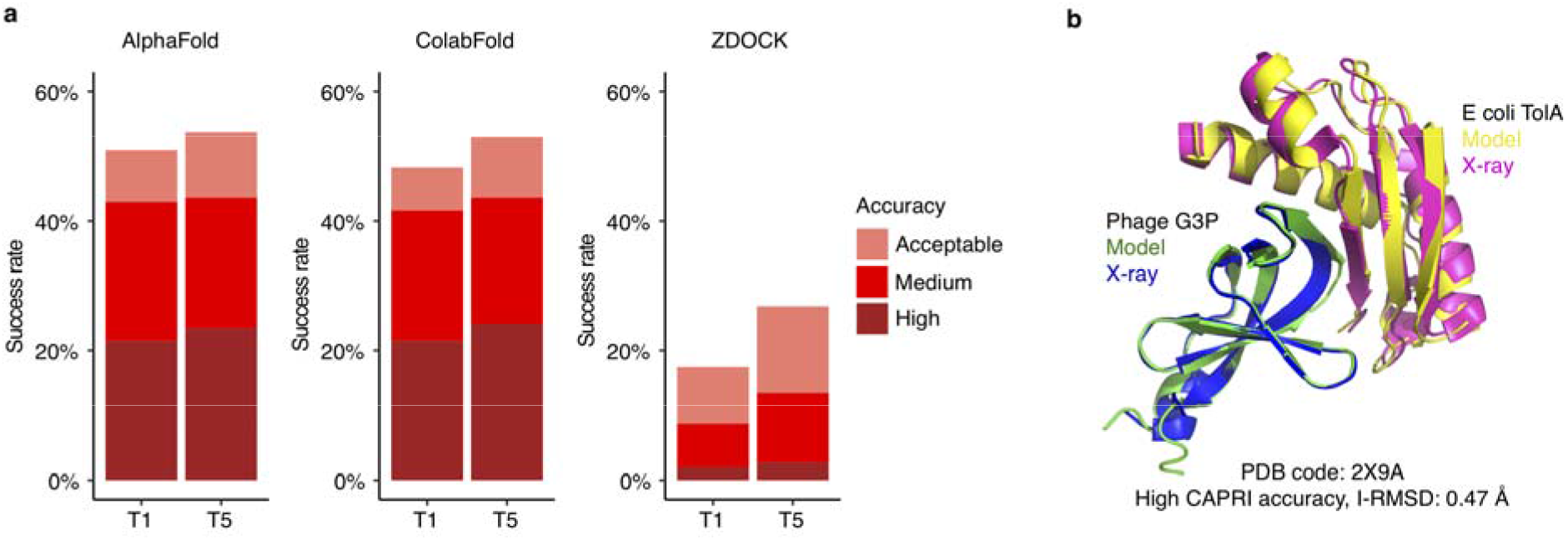
Transient protein-protein complex structure prediction success by AlphaFold, ColabFold and ZDOCK. End-to-end modeling using AlphaFold [24] and ColabFold [28] was performed on 152 complex test cases (details in **Figure S1**). AlphaFold failed to generate predictions for three complexes, thus AlphaFold predictions were obtained for 149 complexes; these 149 test cases were used to calculate success rates in this figure. Docking models were also generated with ZDOCK [29], using unbound protein structures as input. All sets of models were assessed for near-native predictions using CAPRI criteria for High, Medium and Acceptable accuracy. (a) Complex prediction success of AlphaFold, ColabFold, and ZDOCK for the top 1 (T1) and top 5 (T5) models considered. AlphaFold and ColabFold models were ranked by AlphaFold pTM scores, and ZDOCK models were ranked by IRAD scores [44]. The percentage success was calculated as the percentage of test cases with a given model accuracy from the top N models considered. Bars are colored according to the CAPRI quality classes. (b) Example of an accurately predicted complex structure (PDB code: 2×9A) by AlphaFold. This model has High accuracy by CAPRI criteria (I-RMSD = 0.47) and has the highest pTM score (pTM = 0.77) of all 5 models generated for this complex. Structures were superposed by Phage G3P, with the model and the X-ray structure chains colored separately as indicated. For clarity, regions modeled by AlphaFold model but unresolved in the X-ray structure are not shown in the figure.

### Determinants of successful performance

To investigate the determinants of successful performance for AlphaFold, we compared performance across subsets of cases divided by various biological and structural properties (**Figure 2**). As expected, previously assigned test case difficulty classifications, which are based on binding conformational change between unbound and bound structures [20, 23], did not markedly impact the success of AlphaFold. However, the performance was reduced for more challenging “Medium” and “Difficult” cases (Figure 2A) for ZDOCK, which unlike AlphaFold, used unbound protein structures as input. In fact, overall success of AlphaFold was somewhat higher for “Difficult” cases than for “Rigid” and “Medium” case categories (over 75% success, versus approximately 50% success for Acceptable or better prediction in the 5 AlphaFold models). Nonetheless, the limited number of cases in the “Difficult” category (N=15) and the lack of improvement in success for highly accurate models (CAPRI High assessment) among Difficult cases make the implications or generalizability of this effect unclear.

**Figure 2.**
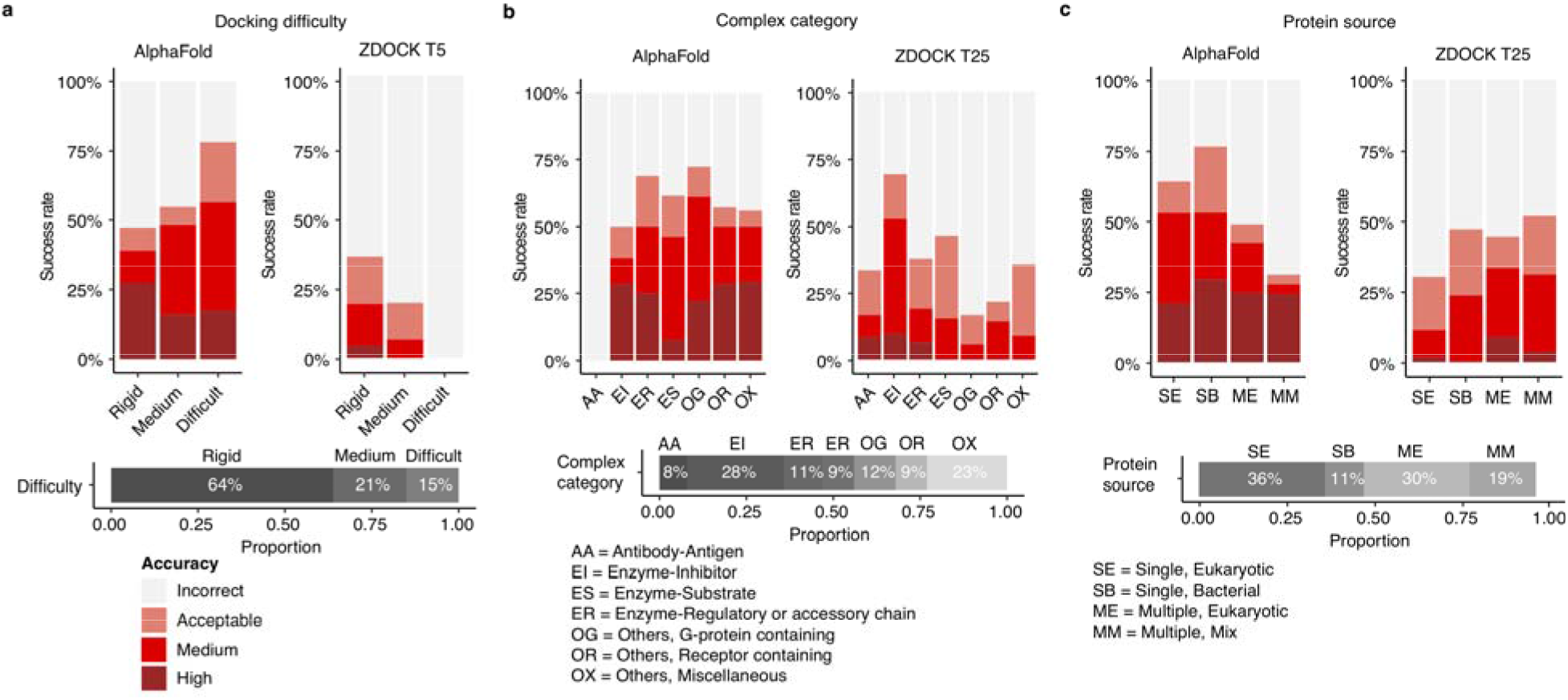
Determinants of successful performance. (a) Prediction success of AlphaFold and ZDOCK, grouped by docking difficulty. Based on binding conformational changes as defined by BM5.5 [20, 23], cases are categorized into “Rigid”, “Medium”, and “Difficult” docking difficulty levels. To evaluate the success rate, all 5 models from AlphaFold and top 5 ZDOCK models were considered. (b) Prediction success of AlphaFold and ZDOCK grouped by complex category. To evaluate the success rate, all 5 models from AlphaFold and top 25 ZDOCK models were considered. The number of ZDOCK models was increased to 25 to allow for sufficient success rates to show its relative performance among the categories. (c) Prediction success of AlphaFold and ZDOCK grouped by protein source organism(s). To evaluate the success rate, all 5 models from AlphaFold and top 25 ZDOCK models were considered. Based on the source of subunit proteins in the complex structures, each case is classified as either “Single, Eukaryotic” (SE, denoting proteins from the same eukaryotic organism), “Single, Bacterial” (SB, denoting proteins from the same bacterial organism), “Multiple, Eukaryotic” (ME, denoting proteins from different eukaryotic organisms), or “Multiple, Mix” (MM, denoting proteins from mixed origins). Two additional protein source classes, corresponding to proteins from different bacterial organisms, and proteins of viral origin, were omitted from the success plot due to limited representation in each category (2 and 3 cases in those classes, respectively). Bars are colored by model accuracy as indicated in (a). The horizontal stacked bars below each success rate plot denote the composition of the categories by class.

Performance across benchmark cases was also assessed by complex category, as well as protein source (**Figure 2b, 2c**). Notably, antibody-antigen complexes had no successfully generated models, while other complex categories considered all showed approximately commensurate levels of AlphaFold performance. There was no major difference observed in AlphaFold success for prediction of complexes with proteins from eukaryotic or bacterial organisms, and while there was a slight reduction in overall success when the two proteins in a complex came from different organisms (which theoretically could impact a cross-interface signal of an MSA), the success for High quality models was approximately the same (∼25%) regardless of single versus multiple source, or organism type. Due to the limited number of virus-containing complex structures (N=3), those cases were not included in percent success calculations.

Other test case properties were investigated for possible relationships with AlphaFold modeling success. Comparison of protein-protein interface sizes (buried surface area; BSA) between cases binned by AlphaFold modeling accuracy (**Figure S2**) did not indicate any impact of interface size on success. We also explored the accuracy of individual chain structural modeling and MSA depth (**Figure S3**); while a range of chain alignment depths (N_eff_) were observed, in most cases the individual ligand and receptor chains were modeled accurately (backbone RMSD with bound component chains < 2.5 Å). Protein complex model accuracies based on interface residue RMSD (I-RMSD) and CAPRI criteria did not show a relationship with minimum chain alignment depth (**Figure S3b**) or maximum subunit chain RMSD (**Figure S3c**), with the latter comparison indicating that incorrect binding mode, versus inaccurate chain folding, was the primary cause of incorrect AlphaFold complex models. AlphaFold models representing incorrect binding mode and inaccurate chain folding are shown in **Figure S3d** (PDB code 1S1Q; Ubiquitin/TSG101 complex) and **Figure S3e** (PDB code 2ABZ, Carboxypeptidase/inhibitor complex).

### Impact of alternative AlphaFold parameters and input

Given the success of AlphaFold with unpaired MSAs, consisting of individual MSAs for each protein, we tested the impact of the use of paired sequences, which represent both chains as a single sequence in the MSA, within the input MSAs. Due to its capability to provide a co-evolution signal between protein residues across an interface, which can then be inferred as cross-interface contacts, use of paired sequences in MSAs has shown promise previously for protein complex structure prediction [32-34]. MSAs with paired sequences were obtained from the ColabFold Google Colab site [28]; these sequence pairs were generated with an automated algorithm intended for prokaryotic proteins, thus a set of 17 cases from BM5.5 was tested that contain two prokaryotic proteins from the same organism. As shown in **Figure S4**, the addition of paired sequences did not appear to improve AlphaFold performance over use of unpaired MSAs as input, while use of paired sequences alone was detrimental to successful complex modeling in some cases. One notable exception was test case 1F6M, which had a relatively high number of paired sequences in the MSA. When paired sequences alone were used, High accuracy models were obtained for test case 1F6M, whereas no hits were obtained when unpaired sequences were included. For comparison, the same paired-only MSAs were input to RoseTTAFold [33], which according to its authors can utilize paired MSAs to predict complex structures; while some accurate models were obtained, we observed lower overall success for the models generated with that method. While it is possible that sub-optimal sequence pairing in the MSAs could have impacted performance of AlphaFold and RoseTTAFold, our limited test here indicates that the already strong performance of AlphaFold using unpaired MSAs does not seem to be markedly affected by addition of paired sequences from a conventional pairing method, and that the absence of the unpaired sequences was detrimental to AlphaFold. We also tested altered parameters for the number of iterative refinement cycles (*N*_*cycle*_) and MSA ensemble size (*N*_*ensemble*_) in AlphaFold, for a subset of the docking test cases selected to represent the antibody, enzyme, and “other” protein complex types, and observed very little effect on predictive performance (**Figure S5**).

### Docking model discrimination by scoring metrics

Given the reported success of AlphaFold in predicting the quality of its monomeric protein models through scores representing local accuracy (pLDDT) and global accuracy (pTM) [24], we tested the discriminative capabilities of these values in the context of protein complex modeling (**Figure 3**). Average pLDDT scores and pTM scores for AlphaFold complex models were both found to discriminate Incorrect versus higher model accuracy classifications, with pTM scores performing moderately better (**Figure 3a**). Comparison of pTM with complex model TM-scores [35] showed a relatively strong correlation of the predicted with the calculated accuracy value (r = 0.82; p < 0.001; **Figure 3b**), while pTM exhibited a significant, though moderately weaker, correlation with I-RMSD of AlphaFold models (r = -0.55, p < 0.001; **Figure 3c**).

**Figure 3.**
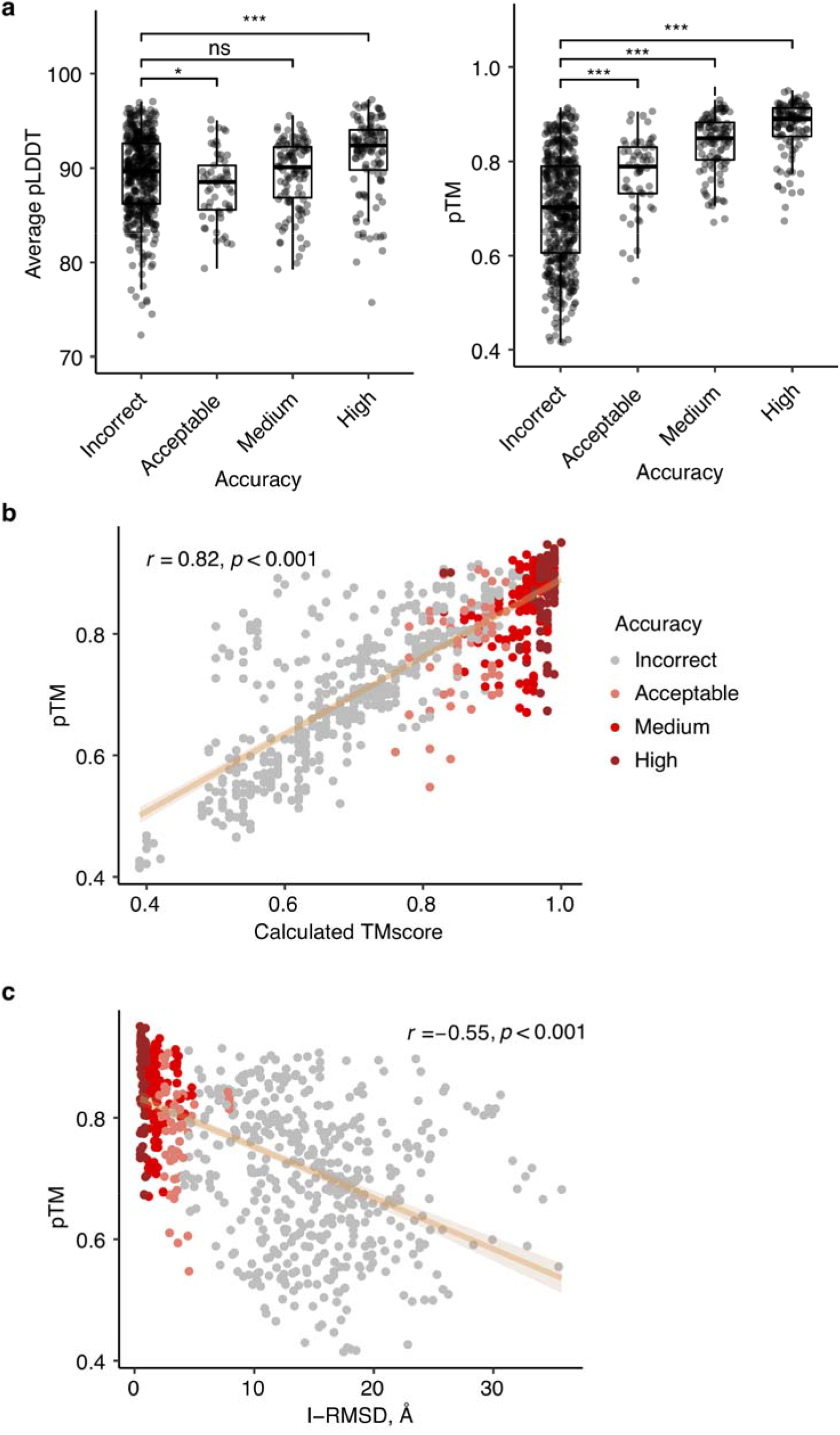
Association between AlphaFold predicted scores and docking model quality. (a) Average pLDDT and pTM per CAPRI criteria. Statistical significance (Wilcoxon rank-sum test) between average pLDDT or pTM of Incorrect vs. Acceptable, Incorrect vs. Medium and Incorrect vs. High CAPRI criteria is indicated at the top (ns: p > 0.05, *: p ≤ 0.05, **: p ≤ 0.01, ***: p ≤ 0.001). (b) Comparisons between pTM and calculated TM score and (c) between pTM and I-RMSD are shown as scatter plots. All 5 models for 149 cases are shown as points, colored by model quality by CAPRI criteria. Linear regression is shown along with the 95% confidence interval (orange area), and Pearson’s correlation coefficients and correlation p-values are denoted in (b) and (c).

While pTM and pLDDT showed some capability to identify correct versus incorrect complex structural models, the overlap in scores between accuracy categories (**Figure 3a**) led us to explore additional scoring functions to predict the structural quality of AlphaFold models (**Figure 4, Table 1**). Given the likely importance of interface residue contacts and packing, versus the folding accuracy of interface-distal protein regions, in discrimination of correct versus incorrect docking models, we tested two residue-level predicted accuracy metrics from AlphaFold (PAE, or Predicted Aligned Error, and pLDDT) for predicted protein-protein interface residues alone, to assess model discrimination capabilities. Alternative formulations of these metrics were tested with more permissive interface definitions, versus the originally tested 4 Å interface cutoff, but no major difference in model assessment accuracy was observed (**Figure S6, Table S2**). Interface PAE and interface pLDDT values showed major improvement compared with average pLDDT and pTM from AlphaFold in discriminating accurate complex models, based on receiver operating characteristic area under the curve (AUC) metrics (**Table 1**), particularly for the discrimination of models in the most populous and divergent Incorrect and High model accuracy categories (AUCs of 0.93 and 0.97 for interface PAE and interface pLDDT, respectively). Relatively high AUC values were also observed for previously reported docking model ranking methods ZRANK2 [15] and IRAD [31], while an interface energy score from Rosetta [36] (Cross-interface binding energy) resulted in the highest model classification accuracy, based on AUC values. While performing lower than the Rosetta binding energy score, some relatively simple protein interface assessments, such as the number of interface hydrogen bonds, showed some capability to classify the accuracy of AlphaFold models.

**Figure 4.**
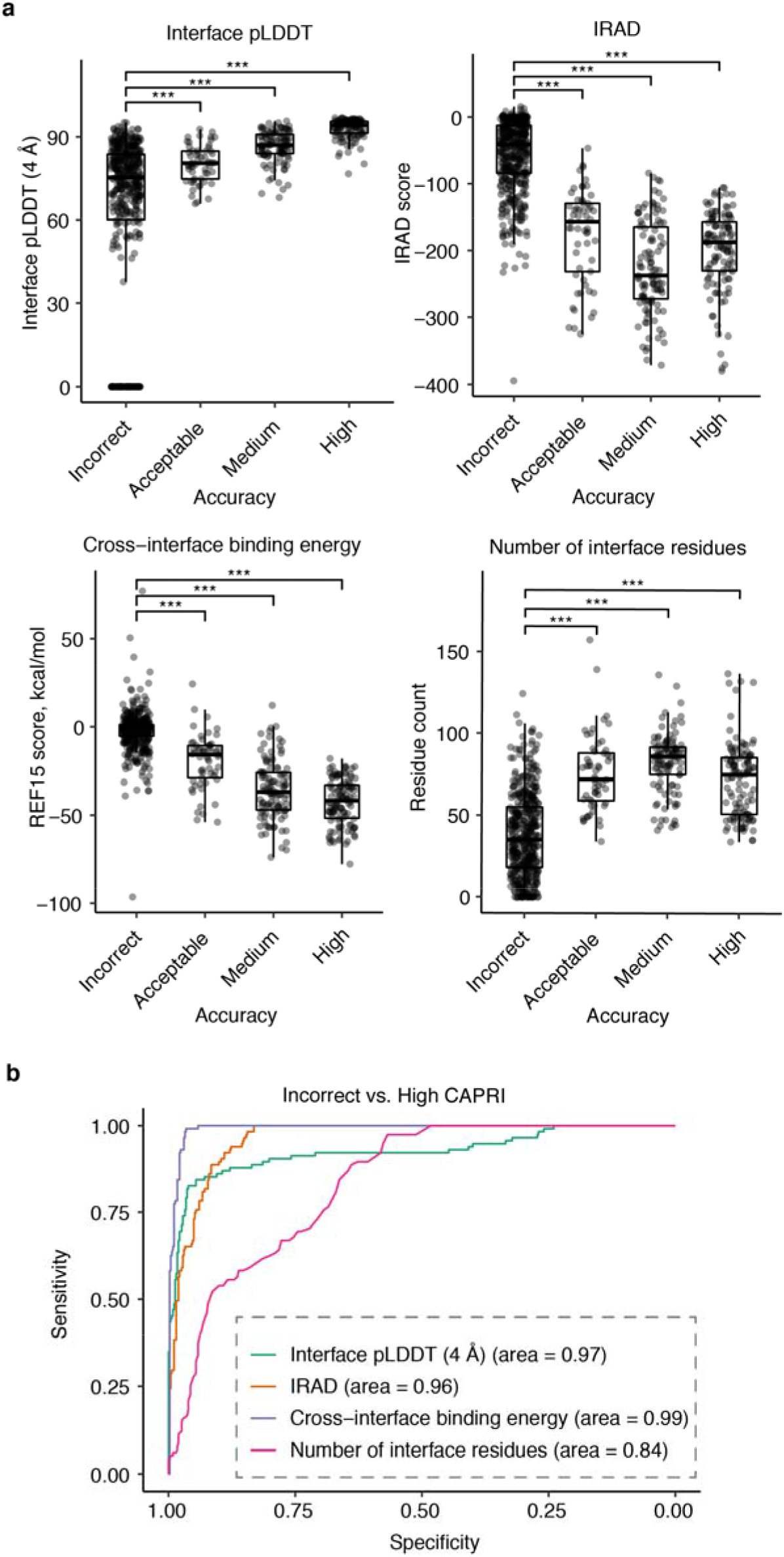
Association between alternative scoring metrics and docking model quality. (a) Distributions of interface pLDDT (4 Å), IRAD, Rosetta cross-interface binding energy, and number of interface residues for AlphaFold models grouped by CAPRI criteria. An interface pLDDT score of 0 was assigned to models without any interface contacts within the distance cutoff (4 Å). Constrained local minimization was performed using Rosetta FastRelax [45] to resolve unfavorable local geometries or clashes in models, and post-relaxation models were scored with IRAD and the Rosetta [36] InterfaceAnalyzer protocol, with the latter used to calculate cross-interface binding energy scores (based on the Rosetta REF15 energy function [47]) and the number of interface residues. Statistical significance values (Wilcoxon rank-sum test) between scores of Incorrect vs. Acceptable, Incorrect vs. Medium, Incorrect vs. High CAPRI criteria are indicated at the top of each plot (***: p ≤ 0.001). Each point corresponds to one AlphaFold model, and all 5 AlphaFold models for 149 test cases are represented. (b) ROC curves among the scoring metrics for classifying Incorrect vs High accuracy models by CAPRI criteria, with corresponding AUC values denoted in parentheses.

**Table 1.** Area under the ROC curve (AUC) values for protein quality classes as a function of scoring metrics.

| Score<sup>a</sup> | Binary classification |  | Multi-class classification |
| --- | --- | --- | --- |
|  | Incorrect vs. High | Incorrect vs. Medium and High |  |
| Average pLDDT | 0.66 | 0.59 | 0.64 |
| Average resolved pLDDT | 0.81 | 0.69 | 0.69 |
| pTM | 0.92 | 0.89 | 0.80 |
| Interface PAE (4 Å) | 0.93 | 0.90 | 0.81 |
| Interface pLDDT (4 Å) | 0.97 | 0.90 | 0.84 |
| IRAD | 0.96 | 0.97 | 0.80 |
| ZRANK | 0.95 | 0.95 | 0.77 |
| Cross-interface binding energy | 0.99 | 0.97 | 0.84 |
| Interface area | 0.90 | 0.92 | 0.77 |
| Number of interface hydrogen bonds | 0.96 | 0.94 | 0.81 |
| Number of interface residues | 0.84 | 0.87 | 0.73 |
| Shape complementarity | 0.91 | 0.85 | 0.79 |

<sup>a</sup>Scoring method. “Average resolved pLDDT”: average pLDDT of the resolved region, “Interface PAE (4 Å)”: average PAE of pairs of interface residues within 4 Å distance cutoff, “Interface pLDDT (4 Å)”: average pLDDT of interface residues within 4 Å distance cutoff. “Cross-interface binding energy”, “Interface area”, “Number of interface hydrogen bonds”, “Number of interface residues” and “Shape complementarity” were calculated using the InterfaceAnalyzer protocol in Rosetta [36].

### Expanded antibody-antigen complex benchmarking

Due to the lack of any successful structural prediction of heterodimeric antibody-antigen complexes from the Benchmark 5.5 set, we assembled a set of 20 additional nonredundant antibody-antigen complexes with known structures to assess AlphaFold accuracy (**Table S3**). These complexes include a variety of antigens, and as the Benchmark 5.5 heterodimer set primarily included nanobodies, a number of single-chain antibodies with both heavy and light chains represented were selected for the additional cases (comprising 17 out of 20 of the cases). While AlphaFold modeling of most of those complex structures resulted in no accurate predictions, surprisingly two of the antibody-antigen complexes were modeled accurately, with Medium CAPRI accuracy models ranked #1 for each complex (**Table S3**, with models shown in **Figure 5a** and **5b**).

**Figure 5.**
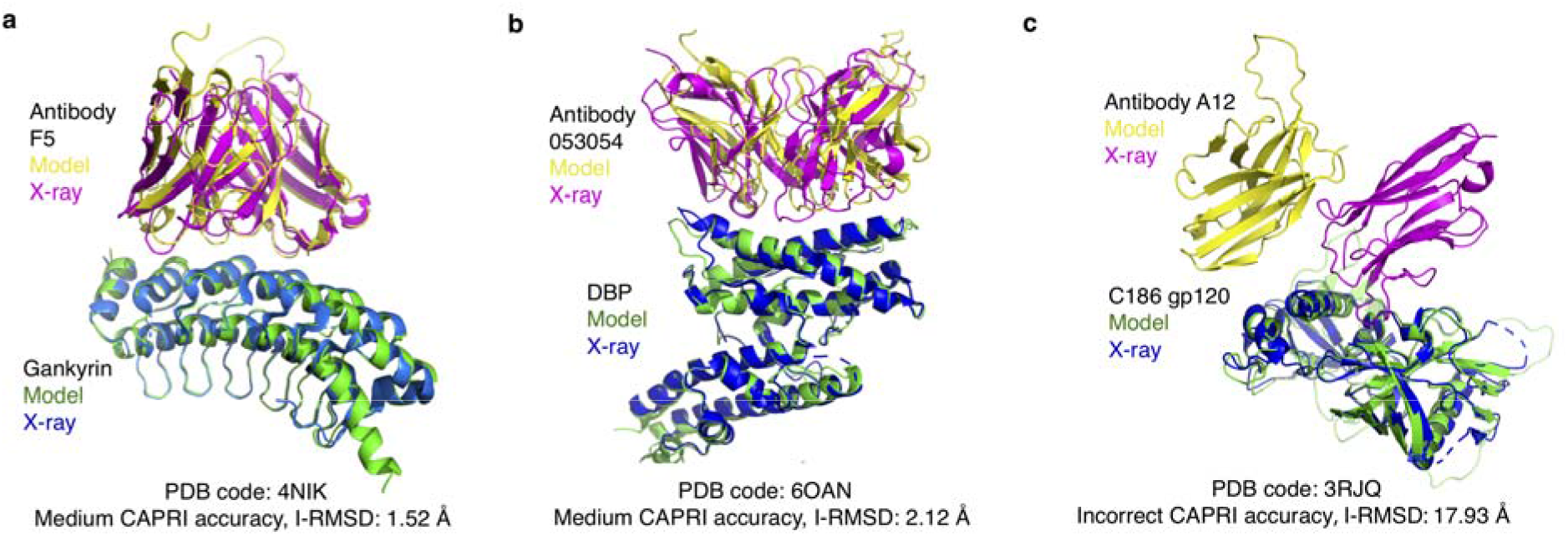
Examples of antibody-antigen complex structure predictions by AlphaFold. (a) Native and top-ranked AlphaFold model (pTM = 0.78) for PDB 4NIK (F5 antibody/human gankyrin complex). This model is of Medium accuracy by CAPRI criteria (I-RMSD = 1.52 Å). Modeled and X-ray complex structures are colored as indicated and shown superposed by gankryin. Unresolved regions modeled by AlphaFold are not shown. (b) Native and top-ranked AlphaFold model (pTM = 0.61) for PDB 6OAN (053054 antibody/P vivax DBP complex). This model is of Medium accuracy by CAPRI criteria (I-RMSD = 2.12 Å). Modeled and X-ray structures are colored as indicated, shown superposed by DBP, and unresolved regions modeled by AlphaFold are not shown. (c) Native and top-ranked AlphaFold model (pTM = 0.66) for PDB 3RJQ (A12 nanobody/HIV C186 gp120 complex), superposed by C186 gp120. This AlphaFold model does not have contacting residues between the proteins within a 5 Å distance cutoff. Structures are colored as indicated in the figure, and unresolved regions modeled by AlphaFold on C186 gp120 are shown in light green.

Inspection of AlphaFold models of antibody-antigen complexes indicated that many of the inaccurate models had few or no contacts between antibody and antigen chains; one example is shown in **Figure 5c**). Indeed, analysis of the percent of cases with no atomic contacts between chains in modeled complexes showed that antibody-antigen cases had relatively high rates of such models in comparison with the other protein complex categories in the benchmark (**Figure S7**).

## Discussion

Our extensive testing of AlphaFold performance on a nonredundant benchmark of protein-protein complexes indicates that AlphaFold is largely successful at predicting binary transient protein complex structures. However, some complexes were not successfully modeled, most notably antibody-antigen interactions, and other categories of test cases showed upper limits of success as well. The two antibody-antigen complex structures that were successfully modeled show that it is possible in principle to perform end-to-end antibody complex modeling with the AlphaFold framework, while many of the incorrect models seem to be readily identifiable based on AlphaFold metrics of predicted interface residues, or previously developed energy-based scoring functions.

AlphaFold’s end-to-end modeling approach represents a major advance and performance improvement over traditional protein-protein docking methods, serving a proof of concept and a possible framework for optimization to specifically model protein-protein interactions with improved accuracy. One initial development in this regard was recently reported by the AlphaFold team (AlphaFold-Multimer) [37], though in accordance with our findings with AlphaFold here, the authors noted that the new version was “generally not able to predict” antibody-antigen complex structures. Another team showed that combination of AlphaFold with a previously developed protein docking method was able to achieve an improvement in docking success [38]. Such developments that improve upon on the complex prediction success of AlphaFold reported here can bring the field closer to solving the longstanding challenge of predictive protein docking.

## Methods

### Protein-protein complex benchmark and additional antibody-antigen test cases

A test set of heterodimeric protein complexes was obtained from Protein-Protein Docking Benchmark 5.5 (BM5.5) [20, 39]. BM5.5 is a set of structures of non-redundant transient protein-protein complexes from the PDB [30], assembled for testing of predictive protein-protein complex modeling algorithms. By filtering for heterodimeric protein-protein complexes in BM5.5, we obtained a total of 152 cases, consisting of 12 antibody-antigen complexes, 72 enzyme-containing complexes and 68 other types of protein complexes. Docking difficulty classifications for test cases were obtained from the BM5.5 site (https://zlab.umassmed.edu/benchmark/), and are based on the extent of binding conformational changes for each complex [20]. Annotations of protein source organisms were obtained from the PDB, and confirmed by manual inspection. Buried surface area for each complex interface was obtained from the BM5.5 site.

Twenty additional antibody-antigen modeling test cases were selected from antibody-antigen complex structures in the SAbDab database [40], screened by resolution (< 3.25 Å) and nonredundancy with any BM5.5 test cases by antibody chain sequences (< 90% antibody variable domain sequence identity) or antigen chain sequence (no hit to antigen chains using default parameters) using the “blastp” executable from the BLAST+ suite [41].

### Complex modeling with AlphaFold

AlphaFold was downloaded from Github (https://github.com/deepmind/alphafold) and installed on a local computing cluster. Sequences of protein chains for the protein-protein complexes were obtained from the PDB “seqres” fasta file and concatenated to form the query sequence for each complex modeling job. Raw MSAs were prepared for each chain with the downloaded published AlphaFold pipeline [24], querying the full databases (UniRef90 version 2020_01, MGnify version 2018_12, Uniclust30 version 2018_08 and BFD). The resulting raw MSAs of the interacting chains were subsequently combined to form the unpaired MSA inputs for complex structure prediction. To generate MSA lines of the same length, gaps equal to the length of the interacting chain were added before or after each sequence. To avoid implicitly biasing the complex structure predictions with knowledge of individual bound protein chain conformations, the use of structural templates was disabled in this study.

To introduce chain breaks, a residue index shift of 200 was added to the junction of interacting chains, as recently implemented in ColabFold [28]. Following the published AlphaFold pipeline, AlphaFold generated five models for each complex, which were ranked in this study by pTM score (predicted TM-score, which is a measure of predicted structure accuracy generated by AlphaFold). After structural predictions were generated, model relaxation by the Amber program [42], which as reported by Jumper et al. [24] was used to ameliorate minor structural defects without impacting accuracy, was replaced by the constrained FastRelax protocol in the Rosetta program [36], as detailed below. To test the impact of varying the ensembling (*N*_*ensemble*_) and recycling (*N*_*cycle*_) parameters on complex modeling accuracy, we increased *N*_*ensemble*_ or *N*_*cycle*_ by modifying those parameters in the AlphaFold Python code, while keeping the input MSAs and sequences/features the same.

Three out of the 152 test cases failed to complete in the AlphaFold pipeline due to GPU memory limits during structure prediction, or errors during feature preparation: 1ZM4, 2OZA, and 1B6C. AlphaFold structure prediction runs were performed on an NVIDIA Titan RTX or NVIDIA Quadro 6000 GPU.

### Complex modeling with ColabFold and RoseTTAFold

Protein-protein complex predictions were generated in ColabFold [28] using its “advanced” interface on Google Colab (https://colab.research.google.com/github/sokrypton/ColabFold/blob/main/beta/AlphaFold_advanced.ipynb). Input protein sequences were identical to those used for AlphaFold modeling. The MMseqs2 method [43] was selected on ColabFold to generate the MSAs, and Amber relaxation of models was disabled. The unpaired MSA predictions are generated between August 20th and August 24th, 2021.

ColabFold enables users to pair alignments for different protein sequences based on UniProt accession numbers; this is a selectable option on the ColabFold interface [28]. Since the protocol is designed to pair prokaryotic protein sequences, MSA pairing was only performed on a subset of cases where both protein chains of the heterodimer complex come from the same prokaryotic organism. Prefiltering of MSAs was enabled prior to pairing, with the minimum coverage with query of 50% and minimum sequence identity with query of 20%. Structural predictions based on paired MSAs were generated on September 4, 2021. The resulting paired MSAs were also used as input to RoseTTAFold [33] on the Robetta web server (https://robetta.bakerlab.org/submit.php) to generate complex models.

All models generated with AlphaFold, ColabFold and RoseTTAFold are made available to the public at: https://piercelab.ibbr.umd.edu/af_complex_models.html.

### Docking model generation with ZDOCK

To enable comparison against a rigid-body docking algorithm, we generated docking models using ZDOCK version 3.0.2 [29]. Unbound protein structures from BM5.5, with HETATMs removed, were used as inputs to ZDOCK. Dense rotational sampling was used, generating 54000 predictions per case. The integration of residue- and atom-based potentials for docking (IRAD) [44] scoring function was used to rank the ZDOCK output models.

### Docking model accuracy assessment

Docking models were assessed using the Critical Assessment of Predicted Interactions (CAPRI) criteria [22] using custom scripts. Based on the structural similarity between docking models and native structures, docking models were classified into four accuracy classes: “High”, “Medium”, “Acceptable” and “Incorrect”. Such structural similarity is assessed by a combination of interface RMSD (I-RMSD), ligand RMSD (L-RMSD), and fraction of native interface residue contacts (fnat). Backbone atoms were used in the I-RMSD and L-RMSD calculations.

### Interface pLDDT and interface PAE calculation

To calculate the interface pLDDT, we averaged the per-residue pLDDT of interface residues. Interface residues are defined as residues with atomic contacts across the interface within the specified distance cutoff. Interface PAE was calculated by averaging the PAE of cross-interface residue pairs with atomic contacts within a given distance cutoff. The interface distance cutoffs tested range from 4 Å to 10 Å. An interface pLDDT score of 0 and an interface PAE score of 35 was assigned to models without any interface contacts within the distance cutoff.

### Structure relaxation using Rosetta

To resolve possible unfavorable geometries or clashes in AlphaFold models, the Rosetta FastRelax protocol [45] was applied to the predicted structures prior to scoring of the models using interface analysis protocols (IRAD, ZRANK2, and Rosetta). Parameter flags used in FastRelax (“relax” executable in Rosetta 3.12 [36]) are:

- relax:constrain_relax_to_start_coords
- relax:coord_constrain_sidechains
- relax:ramp_constraints false
- ex1
- ex2
- use_input_sc
- no_optH false
- flip_HNQ
- nstruct 1

### Docking model scoring with IRAD, ZRANK2, and Rosetta InterfaceAnalyzer

Post-relax model structures were used as inputs to obtain IRAD [31], ZRANK2 [15] and Rosetta InterfaceAnalyzer [46] scores. IRAD and ZRANK2 scores were obtained from the downloaded “irad” executable program. InterfaceAnalyzer scores were obtained using the. “InterfaceAnalyzer” executable in Rosetta v. 3.12, with default parameters; the InterfaceAnalyzer protocol computes and outputs interface energetic scores using the Rosetta REF15 function [47], along with REF15 component terms and other interface structure metrics.

### Number of effective sequences

The number of effective sequences (N_eff_) is used as a measure of the MSA depth. N_eff_ score is defined as the number of clusters after the raw MSA inputs were clustered at the 62 % sequence identity using CD-HIT [48] with the word length of 4, as used previously [49].

### TM-score calculations

TM-scores were calculated using TM-score executable [35] by comparing the structural similarity between experimentally determined structures and AlphaFold models. Residues that were unresolved in experimentally determined structures were removed from AlphaFold models before the calculation of TM-scores.

### Figures, statistical analysis, and AUC calculations

Figures of structures were generated using PyMOL version 2.4 (Schrodinger, Inc.). Box plots, line plots and bar plots were generated with the ggplot2 package [50] in R (r-project.org), and heatmaps were generated with the ComplexHeatmap package [51] in R. Pearson correlations and their p-values were calculated with ggpubr package in R. Wilcoxon rank-sum test was performed using ggsignif package in R. Binary and multi-class ROC curves with AUC values were calculated with the pROC and multiROC packages, respectively, in R.

## Acknowledgements

We thank the AlphaFold, ColabFold and RoseTTAFold teams for developing and distributing the structural prediction algorithms that enabled this study. We are grateful to Ipsa Mittra (University of Maryland) for assistance with generating structural predictions with ColabFold. We are additionally thankful to Johnathan Guest, Ragul Gowthaman and John Moult (University of Maryland Institute for Bioscience and Biotechnology Research) for helpful discussions and suggestions. The Institute for Bioscience and Biotechnology Research computing facility and staff, including Thomas Lane and Christian Presley, provided resources and assistance with installing and running AlphaFold. This work was supported by startup funding from the University of Maryland (B.G.P.) and NIH R01 GM126299 (B.G.P.).

