## Supplemental Figures and Tables for "Benchmarking AlphaFold for protein complex modeling reveals accuracy determinants"

**Supplementary Figure Legends**

**Figure S1. Predictive modeling success of AlphaFold, ColabFold, and ZDOCK.** Cases are grouped and sorted by complex type. For reference, complex category, buried surface area (BSA, $Å^2$) and docking difficulty based on binding conformational changes are indicated on the left. Complex category: Enzyme-inhibitor (EI), enzyme complex with a regulatory or accessory chain (ER), enzyme-substrate (ES), others, G-protein containing (OG), others, receptor-containing (OR); others, miscellaneous (OX), antibody-antigen (AA). Success for top 1 and top 5 (T1, T5) ranked predictions is shown, colored by CAPRI model accuracy as indicated in the key on right (Hits). Three cases failed to complete in AlphaFold and are shown as dark gray cells ("ND" under Hits).

**Figure S2. Comparison of binding interface size and docking success.** Distributions of interface size as measured by buried surface area (BSA, $Å^2$) are shown, grouped by model accuracy according to CAPRI criteria for AlphaFold top 5 (left) or ZDOCK top 25 (right) models. BSA values were calculated as the change in solvent accessible area upon binding, based on the X-ray structures of the complexes, and were obtained from the BM5.5 site. Statistical significance (Wilcoxon rank-sum test) between BSA values of cases with Incorrect vs. High CAPRI accuracy models is indicated at the top of each plot (ns: $p > 0.5$).

**Figure S3. Comparison of MSA depth, subunit accuracy, and AlphaFold model quality.** (a) Scatter plots depicting the relationship between ligand or receptor intersect RMSD and MSA depth (log scale; measured by $N_{eff}$, see Materials and Methods for details). Each point represents the top pTM model from each of the 149 cases modeled by AlphaFold. Points are colored by I-RMSD of the docking model. Scatter plot depicting the relationship between the docking model I-RMSD and (b) minimum ligand and receptor MSA depth (log scale, measured by $N_{eff}$), or (c) Maximum subunit (ligand or receptor) RMSD from X-ray structure. Points are colored by CAPRI accuracy and represent the top-ranked model (by pTM score) from each of the 149 cases modeled by AlphaFold. (d) Native and top-ranked AlphaFold model (pTM = 0.89) for PDB 1S1Q, superposed by TSG101. (e) Native and top-ranked AlphaFold model (pTM = 0.81) for PDB 2ABZ, superposed by Carboxypeptidase A1.

Model and the X-ray structure chains are colored separately as indicated. Unresolved regions modeled by AlphaFold were omitted from the figures.

**Figure S4. The impact of MSA pairing on prediction accuracy.** MSAs were generated using MMseqs2 using the "advanced" interface of ColabFold [1]. Pairing was performed in ColabFold on a total of 17 cases whose ligand and receptor proteins come from the same prokaryotic organism. Cases in the heatmap were sorted by the paired MSA depth ($N_{eff}$; see Material and Methods for details) from the largest to the smallest values. Structural predictions were generated with the "advanced" interface of ColabFold, and RoseTTAFold [2] (through the Robetta server). All models were assessed for near-native predictions within the top-ranked (T1) and top 5 (T5) models using CAPRI criteria. Complex category: Enzyme-inhibitor (EI), enzyme complex with a regulatory or accessory chain (ER), enzyme-substrate (ES), others, receptor-containing (OR); others, miscellaneous (OX).

**Figure S5. Testing the impact of alternative parameters on AlphaFold accuracy.** Models were generated with AlphaFold for 21 test cases, representing three cases from each of the 7 complex categories, chosen by random. By default, $N_{cycle}$ is set to 3 and $N_{ensemble}$ is set to 1. To test the impact of using alternative parameters, $N_{cycle}$ and $N_{ensemble}$ were respectively increased to 48 and 8, while all other components of AlphaFold pipeline were kept constant. All models were assessed for near-native predictions using CAPRI criteria for High, Medium and Acceptable accuracy in the top-ranked (T1) and top 5 (T5) models. Complex category: Enzyme-inhibitor (EI), enzyme complex with a regulatory or accessory chain (ER), enzyme-substrate (ES), others, G-protein containing (OG), others, receptor-containing (OR); others, miscellaneous (OX), antibody-antigen (AA).

**Figure S6. Alternative formulation of interface accuracy metrics calculated from AlphaFold predictions (PAE, pLDDT)** (a) Interface PAE within a distance cutoff of 4 Å, 5 Å, 6 Å, 7 Å and 10 Å grouped by docking model accuracy. (b) Interface pLDDT within a distance cutoff of 4 Å, 5 Å, 6 Å, 7 Å and 10 Å, grouped by docking model accuracy. An interface pLDDT score of 0 and an interface PAE score of 35 are assigned to models without interface contacts within the distance cutoff specified. See also Table S2 for AUC values for interface pLDDT and interface PAE as classifiers for docking model accuracy.

**Figure S7**. **Distribution of model predictions without inter-chain atomic contacts.** The bars denote the percentage of incorrectly predicted cases with at least one model with no interface contacts within 5 Å distance cutoff per complex category. Complex category: Antibody-antigen (AA), enzyme-inhibitor (EI), enzyme complex with a regulatory or accessory chain (ER), enzyme-substrate (ES), others, G-protein containing (OG), others, receptor-containing (OR); others, miscellaneous (OX).

### Figure S1

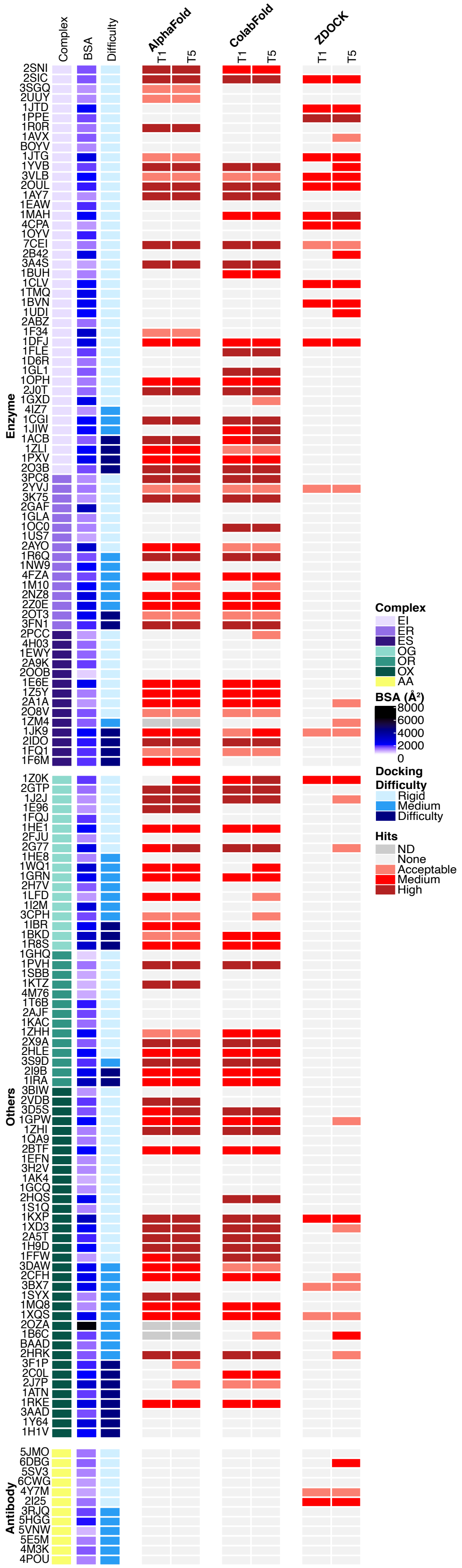
Complex
BSA
Difficulty
AlphaFold
ColabFold
ZDOCK
T1
T5
Enzyme
2SNI
2SIC
3SGQ
2UUY
1JTD
1PPE
1R0R
1AVX
BOYV
1JTG
1YVB
3VLB
2OUL
1AY7
1EAW
1MAH
4CPA
1OYV
7CEI
2B42
3A4S
1BUH
1CLV
1TMQ
1BVN
1UDI
2ABZ
1F34
1DFJ
1FLE
1D6R
1GL1
1OPH
2J0T
1GXD
4IZ7
1CGI
1JIW
1ACB
1ZLI
1PXV
2O3B
3PC8
2YVJ
3K75
2GAF
1GLA
1OC0
1US7
2AYO
1R6Q
1NW9
4FZA
1M10
2NZ8
2Z0E
2OT3
3FN1
2PCC
4H03
1EWY
2A9K
2OOB
1E6E
1Z5Y
2A1A
2O8V
1ZM4
1JK9
2IDO
1FQ1
1F6M
Others
1Z0K
2GTP
1J2J
1E96
1FQJ
1HE1
2FJU
2G77
1HE8
1WQ1
1GRN
2H7V
1LFD
1I2M
3CPH
1IBR
1BKD
1R8S
1GHQ
1PVH
1SBB
1KTZ
4M76
1T6B
2AJF
1KAC
1ZHH
2X9A
2HLE
3S9D
2I9B
1IRA
3BIW
2VDB
3D5S
1GPW
1ZHI
1QA9
2BTF
1EFN
3H2V
1AK4
1GCQ
2HQS
1S1Q
1KXP
1XD3
2A5T
1H9D
1FFW
3DAW
2CFH
3BX7
1SYX
1MQ8
1XQS
2OZA
1B6C
BAAD
2HRK
3F1P
2C0L
2J7P
1ATN
1RKE
3AAD
1Y64
1H1V
Antibody
5JMO
6DBG
5SV3
6CWG
4Y7M
2I25
3RJQ
5HGG
5VNW
5E5M
4M3K
4POU
Complex
EI
ER
ES
OG
OR
OX
AA
BSA (Å²)
8000
6000
4000
2000
0
Docking Difficulty
Rigid
Medium
Difficulty
Hits
ND
None
Acceptable
Medium
High

**Figure S2**

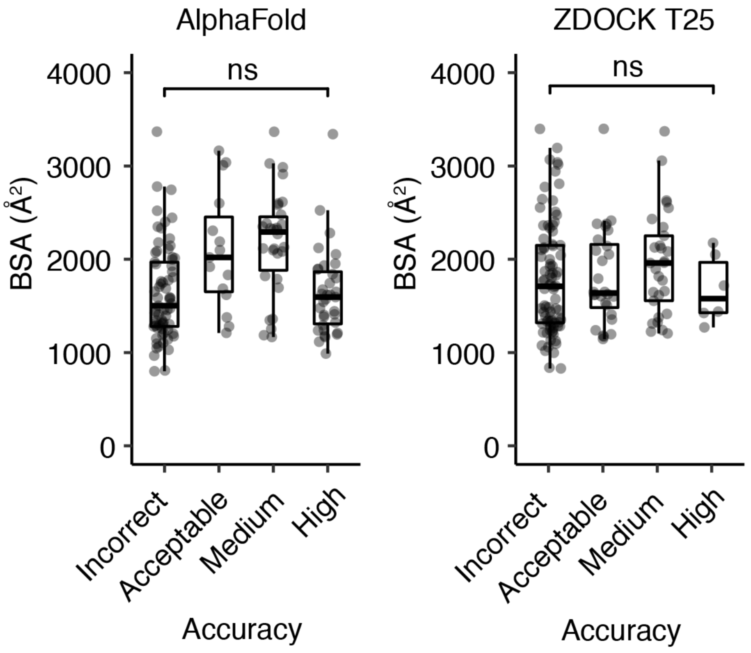

AlphaFold
ZDOCK T25
ns
ns
BSA (Å²)
BSA (Å²)
4000
3000
2000
1000
0
Incorrect
Acceptable
Medium
High
Accuracy
Accuracy

**Figure S3**

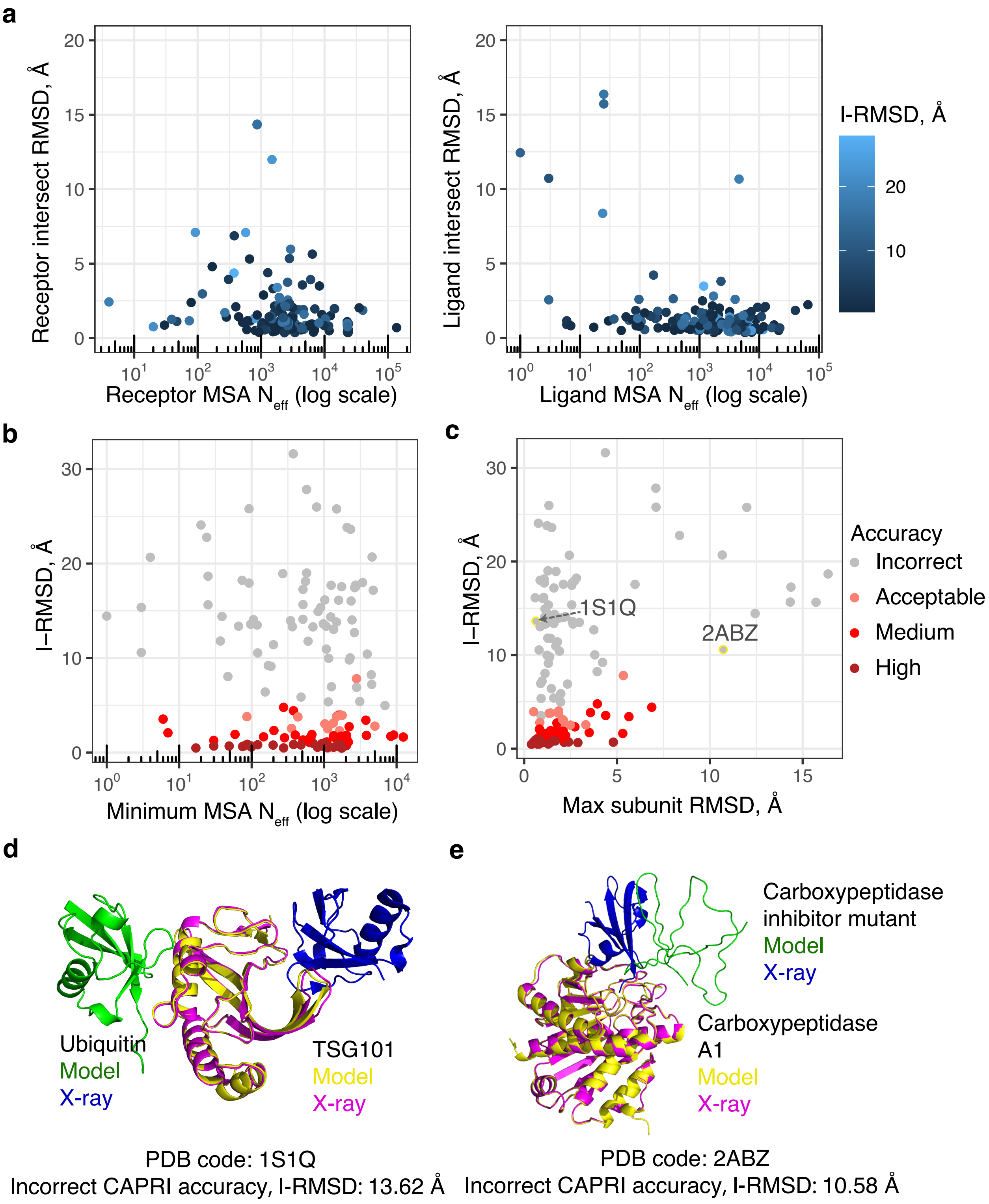

a
Receptor intersect RMSD, Å
Receptor MSA $N_{eff}$ (log scale)
Ligand intersect RMSD, Å
Ligand MSA $N_{eff}$ (log scale)
I-RMSD, Å
b
I–RMSD, Å
Minimum MSA $N_{eff}$ (log scale)
c
Max subunit RMSD, Å
1S1Q
2ABZ
Accuracy
Incorrect
Acceptable
Medium
High
d
Ubiquitin
Model
X-ray
TSG101
Model
X-ray
PDB code: 1S1Q
Incorrect CAPRI accuracy, I-RMSD: 13.62 Å
e
Carboxypeptidase inhibitor mutant
Model
X-ray
Carboxypeptidase A1
Model
X-ray
PDB code: 2ABZ
Incorrect CAPRI accuracy, I-RMSD: 10.58 Å

**Figure S4**

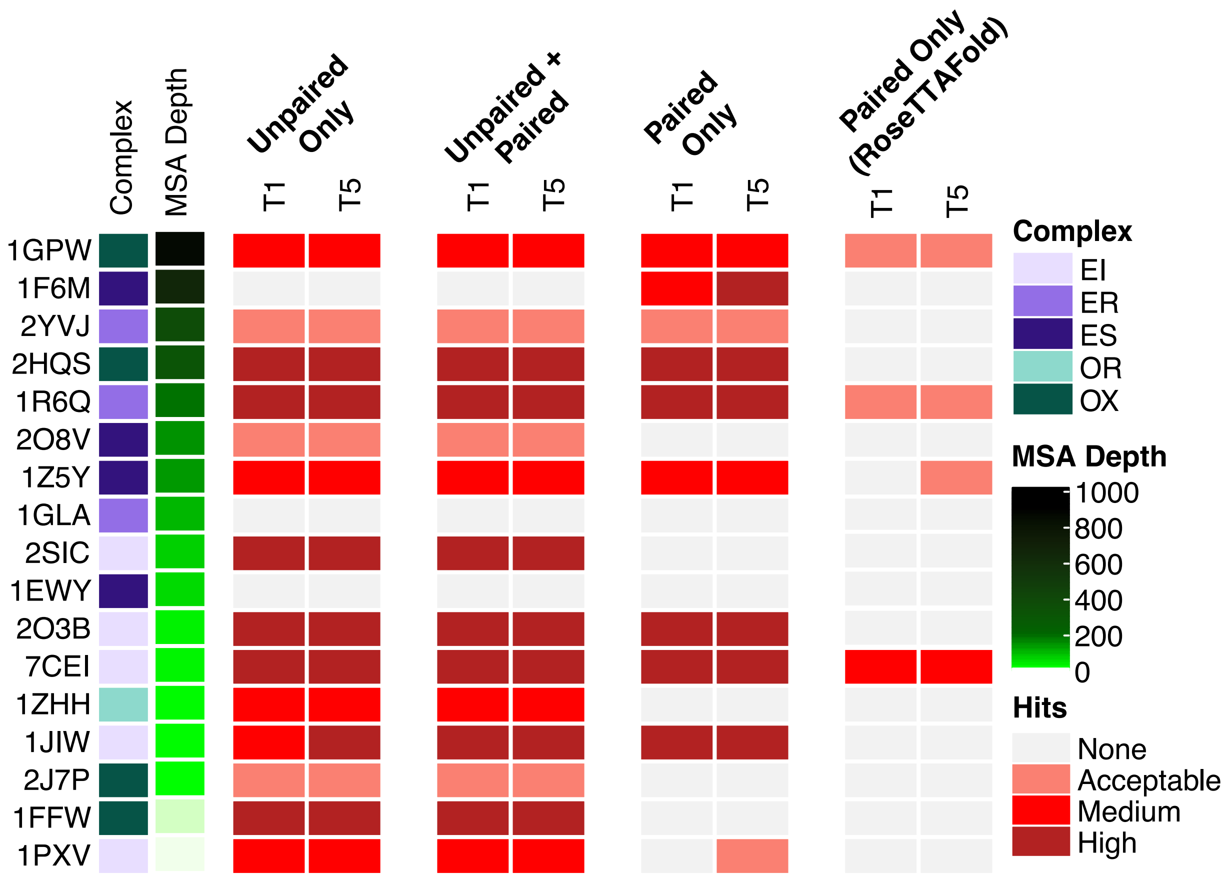
Complex
MSA Depth
Unpaired Only
Unpaired + Paired
Paired Only
Paired Only (RoseTTAFold)
T1
T5
1GPW
1F6M
2YVJ
2HQS
1R6Q
2O8V
1Z5Y
1GLA
2SIC
1EWY
2O3B
7CEI
1ZHH
1JIW
2J7P
1FFW
1PXV
Complex
EI
ER
ES
OR
OX
MSA Depth
1000
800
600
400
200
0
Hits
None
Acceptable
Medium
High

**Figure S5**

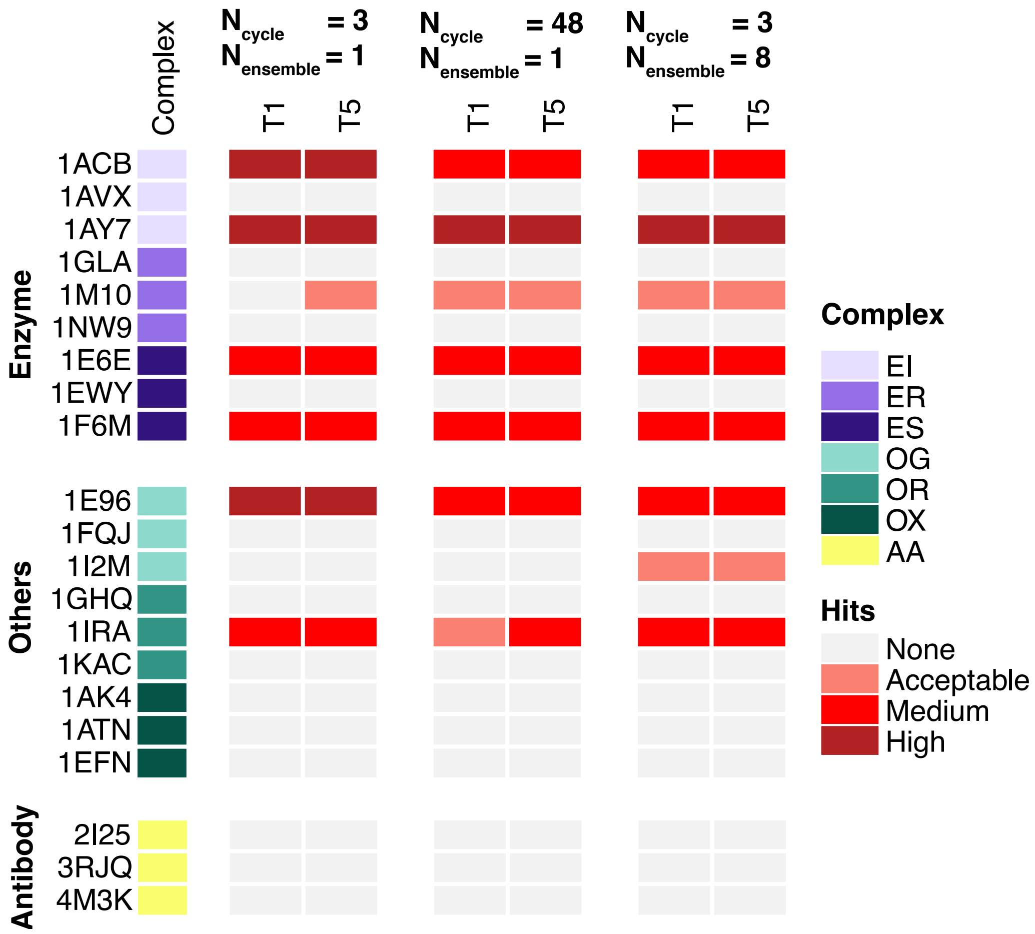

Complex
N cycle = 3
N ensemble = 1
N cycle = 48
N ensemble = 1
N cycle = 3
N ensemble = 8
T1
T5
T1
T5
T1
T5
Enzyme
1ACB
1AVX
1AY7
1GLA
1M10
1NW9
1E6E
1EWY
1F6M
Others
1E96
1FQJ
1I2M
1GHQ
1IRA
1KAC
1AK4
1ATN
1EFN
Antibody
2I25
3RJQ
4M3K
Complex
EI
ER
ES
OG
OR
OX
AA
Hits
None
Acceptable
Medium
High

### Figure S6

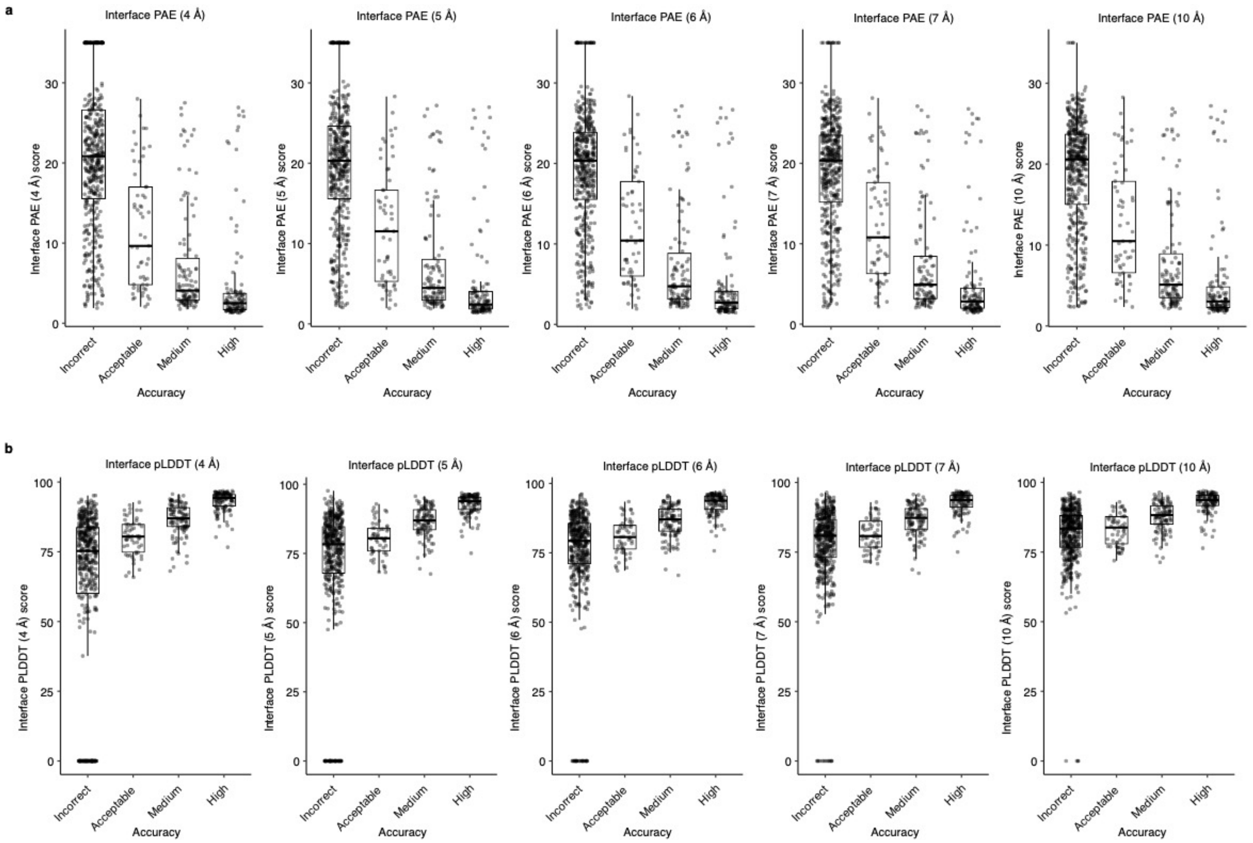
a
Interface PAE (4 Å)
Interface PAE (5 Å)
Interface PAE (6 Å)
Interface PAE (7 Å)
Interface PAE (10 Å)
Interface PAE (4 Å) score
Interface PAE (5 Å) score
Interface PAE (6 Å) score
Interface PAE (7 Å) score
Interface PAE (10 Å) score
Incorrect
Acceptable
Medium
High
Accuracy
b
Interface pLDDT (4 Å)
Interface pLDDT (5 Å)
Interface pLDDT (6 Å)
Interface pLDDT (7 Å)
Interface pLDDT (10 Å)
Interface PLDDT (4 Å) score
Interface PLDDT (5 Å) score
Interface PLDDT (6 Å) score
Interface PLDDT (7 Å) score
Interface PLDDT (10 Å) score
Incorrect
Acceptable
Medium
High
Accuracy

**Figure S7**

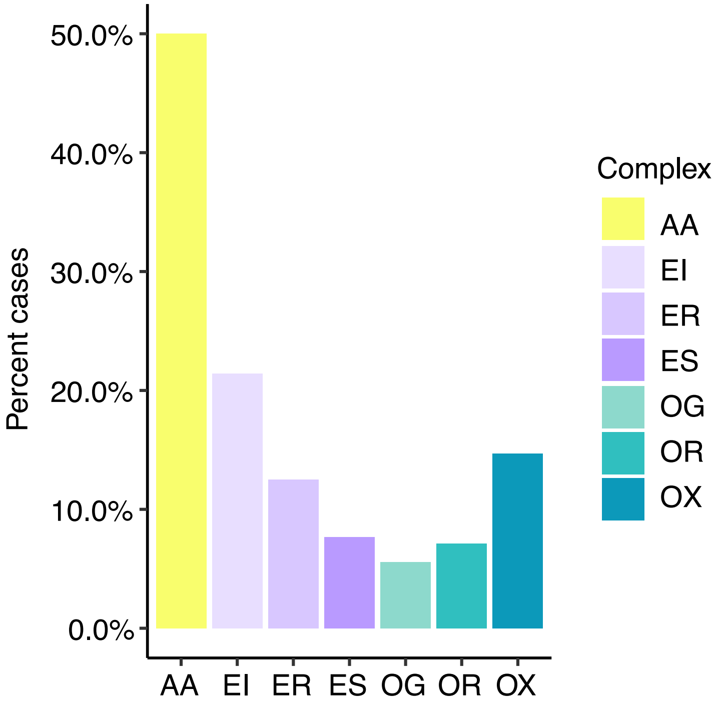

50.0%
40.0%
30.0%
20.0%
10.0%
0.0%
Percent cases
AA
EI
ER
ES
OG
OR
OX
Complex
AA
EI
ER
ES
OG
OR
OX

**Table S1.** Heterodimeric protein-protein complexes from BM5.5 tested in this study.

| PDB code | Receptor | Ligand | Complex Category[a] | BSA (Å²)[b] | Protein sources[c] |
|---|---|---|---|---|---|
| 1ACB | Chymotrypsin | Eglin C | EI | 1544 | BOS TAURUS; HIRUDO MEDICINALIS |
| 1AK4 | Cyclophilin | HIV capsid | OX | 1029 | HOMO SAPIENS; HUMAN IMMUNODEFICIENCY VIRUS 1 |
| 1ATN | Actin | Dnase I | OX | 1774 | ORYCTOLAGUS CUNICULUS; BOS TAURUS |
| 1AVX | Porcine trypsin | Soybean trypsin inhibitor | EI | 1585 | SUS SCROFA; GLYCINE MAX |
| 1AY7 | RNase Sa | Barstar | EI | 1237 | STREPTOMYCES AUREOFACIENS; BACILLUS AMYLOLIQUEFACIENS |
| 1B6C | FKBP binding protein | TGFbeta receptor | OX | 1752 | HOMO SAPIENS |
| 1BKD | Ras GTPase | Son of sevenless | OG | 3163 | HOMO SAPIENS |
| 1BUH | CDK2 kinase | Ckshs1 | EI | 1324 | HOMO SAPIENS |
| 1BVN | Alpha-amylase | Tendamistat | EI | 2222 | SUS SCROFA; STREPTOMYCES TENDAE |
| 1CGI | Bovine chymotrypsinogen | PSTI | EI | 2053 | BOS TAURUS; HOMO SAPIENS |
| 1CLV | Alpha-amylase | Alpha-amylase inhibitor | EI | 2087 | TENEBRIO MOLITOR; SYNTHETIC CONSTRUCT |
| 1D6R | Bovine trypsin | Bowman-Birk inhibitor | EI | 1408 | BOS TAURUS; GLYCINE MAX |
| 1DFJ | Ribonuclease A | Rnase inhibitor | EI | 2582 | BOS TAURUS; SUS SCROFA |
| 1E6E | Adrenoxin reductase | Adrenoxin | ES | 2315 | BOS TAURUS |
| 1E96 | Rac GTPase | p67 Phox | OG | 1179 | HOMO SAPIENS |
| 1EAW | Matriptase | BPTI | EI | 1866 | HOMO SAPIENS; BOS TAURUS |
| 1EFN | HIV-1-NEF protein | SH3 domain | OX | 1254 | HOMO SAPIENS; HUMAN IMMUNODEFICIENCY VIRUS 1 |
| 1EWY | Ferredoxin reductase | Ferredoxin | ES | 1502 | NOSTOC SP. PCC 7119 |
| 1F34 | Porcine pepsin | Ascaris inhibitor 3 | EI | 3038 | ASCARIS SUUM; SUS SCROFA |
| 1F6M | Thioredoxin reductse | Thioredoxin 1 | ES | 1821 | ESCHERICHIA COLI |
| 1FFW | Chemotaxis protein CheY | Chemotaxis protein CheA | OX | 1166 | ESCHERICHIA COLI |
| 1FLE | Elastase | Elafin | EI | 1763 | HOMO SAPIENS; SUS SCROFA |
| 1FQ1 | CDK2 kinase | CDK inhibitor 3 | ES | 1832 | HOMO SAPIENS |
| 1FQJ | Gt-alpha | RGS9 | OG | 1806 | BOS TAURUS; RATTUS NORVEGICUS |
| 1GCQ | GRB2 C-ter SH3 domain | Vav N-ter SH3 domain | OX | 1208 | HOMO SAPIENS; MUS MUSCULUS |
| 1GHQ | Complement C3 | Epstein-Barr virus receptor CR2 | OR | 800 | HOMO SAPIENS |
| 1GL1 | Alpha-chymotrypsin | Protease inhibitor LCMI II | EI | 1591 | BOS TAURUS; SYNTHETIC CONSTRUCT |
| 1GLA | Glycerol Kinase | Glucose specific phosphocarrier | ER | 1304 | ESCHERICHIA COLI |
| 1GPW | HISF protein | Amidotransferase HISH | OX | 2097 | THERMOTOGA MARITIMA |
| 1GRN | CDC42 GTPase | CDC42 GAP | OG | 2332 | HOMO SAPIENS |
| 1GXD | proMMP2 type IV collagenase | Metalloproteinase inhibitor 2 | EI | 2445 | HOMO SAPIENS |
| 1H1V | Actin | Gelsolin | OX | 2071 | HOMO SAPIENS; ORYCTOLAGUS CUNICULUS |
| 1H9D | Runx1 domain of CBFAlpha1 | Dimerisation domain of CBF-Beta | OX | 2121 | HOMO SAPIENS; SYNTHETIC CONSTRUCT |

| | | | | | |
|---|---|---|---|---|---|
| 1HE1 | Rac GTPase | Pseudomonas toxin GAP dom. | OG | 2113 | PSEUDOMONAS AERUGINOSA; HOMO SAPIENS |
| 1HE8 | Ras GTPase | PIP3 kinase | OG | 1305 | HOMO SAPIENS |
| 1I2M | Ran GTPase | RCC1 | OG | 2779 | HOMO SAPIENS |
| 1IBR | Ran GTPase | Importin beta | OG | 2270 | HOMO SAPIENS |
| 1IRA | Interleukin-1 receptor | Interleukin-1 receptor antagonist protein | OR | 3367 | HOMO SAPIENS |
| 1J2J | Arf1 GTPase | GAT domain of GGA1 | OG | 1209 | MUS MUSCULUS; HOMO SAPIENS |
| 1JIW | Alkaline metalloproteinase | Proteinase inhibitor | EI | 1997 | PSEUDOMONAS AERUGINOSA |
| 1JK9 | CCS metallochaperone | SOD1 superoxide dismutase | ES | 2130 | SACCHAROMYCES CEREVISIAE |
| 1JTD | BLIP-II | TEM-1 beta-lactamase | EI | 2180 | ESCHERICHIA COLI; STREPTOMYCES EXFOLIATUS |
| 1JTG | Beta-lactamase inhibitor protein | Beta-lactamase TEM-1 | EI | 2600 | ESCHERICHIA COLI; STREPTOMYCES CLAVULIGERUS |
| 1KAC | Adenovirus fiber knob protein | Adenovirus receptor | OR | 1456 | HUMAN ADENOVIRUS 12; HOMO SAPIENS |
| 1KTZ | TGF-Beta | TGF-Beta receptor | OR | 989 | HOMO SAPIENS |
| 1KXP | Actin | Vitamin D binding protein | OX | 3341 | ORYCTOLAGUS CUNICULUS; HOMO SAPIENS |
| 1LFD | Ras | RalGDS Ras-interacting domain | OG | 1167 | RATTUS NORVEGICUS; HOMO SAPIENS |
| 1M10 | Von willebrand factor dom. A1 | Glycoprotein IB-Alpha | ER | 2097 | HOMO SAPIENS |
| 1MAH | Acetylcholinesterase | Fasciculin | EI | 2145 | MUS MUSCULUS; DENDROASPIS ANGUSTICEPS |
| 1MQ8 | ICAM-1 domain 1-2 | Integrin Alpha-L I domain | OX | 1253 | HOMO SAPIENS |
| 1NW9 | Capase-9 | BIR3-XIAP | ER | 2112 | HOMO SAPIENS |
| 1OC0 | Plasminogen activator inhibitor-1 | Vitronectin Somatomedin B domain | ER | 1313 | HOMO SAPIENS |
| 1OPH | Alpha-1-antitrypsin | Trypsinogen | EI | 1360 | HOMO SAPIENS; BOS TAURUS |
| 1OYV | Subtilisin Carlsberg | Two-headed tomato inhibitor-II | EI | 1930 | BACILLUS LICHENIFORMIS; SOLANUM LYCOPERSICUM |
| 1PPE | Bovine trypsin | CMTI-1 squash inhibitor | EI | 1688 | BOS TAURUS |
| 1PVH | IL6 receptor Beta chain D2-D3 domains | Leukemia inhibitory factor | OR | 1403 | HOMO SAPIENS |
| 1PXV | Cystein protease | Cystein protease inhibitor | EI | 2336 | STAPHYLOCOCCUS AU REUS |
| 1QA9 | CD2 | CD58 | OX | 1353 | HOMO SAPIENS |
| 1R0R | Subtilisin carlsberg | OMTKY | EI | 1409 | BACILLUS LICHENIFORMIS; MELEAGRIS GALLOPAVO |
| 1R6Q | Clp protease subunit ClpA | Clp protease adaptor protein ClpS | ER | 1651 | ESCHERICHIA COLI |
| 1R8S | Arf1 GTPase | Sec 7 domain | OG | 2986 | BOS TAURUS; HOMO SAPIENS |
| 1RKE | Vinculin head | Vinculin tail | OX | 2614 | HOMO SAPIENS |
| 1S1Q | UEV domain | Ubiquitin | OX | 1288 | HOMO SAPIENS |
| 1SBB | T-cell receptor Beta | Staphylococcus enterotoxin B | OR | 1064 | MUS MUSCULUS; STAPHYLOCOCCUS AUREUS |
| 1SYX | Spliceosomal U5 15 kDa protein | CD2 receptor binding protein 2 C-ter fragment | OX | 1293 | HOMO SAPIENS |

| | | | | | |
|---|---|---|---|---|---|
| 1T6B | Anthrax protective antigen | Anthrax toxin receptor | OR | 1948 | BACILLUS ANTHRACIS; HOMO SAPIENS |
| 1TMQ | alpha-amylase | RAGI inhibitor | EI | 2401 | TENEBRIO MOLITOR; ELEUSINE CORACANA |
| 1UDI | Uracyl-DNA glycosylase | Glycosylase inhibitor | EI | 2022 | HERPES SIMPLEX VIRUS (TYPE 1 / STRAIN 17); BACILLUS PHAGE PBS1 |
| 1US7 | Heat shock protein 82 N-ter domain | HSP 90 co-chaperone CDC37 C-ter domain | ER | 1095 | SACCHAROMYCES CEREVISIAE; HOMO SAPIENS |
| 1WQ1 | Ras GTPase | Ras GAP | OG | 2913 | HOMO SAPIENS |
| 1XD3 | UCH-L3 | Ubiquitin | OX | 2281 | HOMO SAPIENS |
| 1XQS | HspBP1 | Hsp70 ATPase domain | OX | 2350 | HOMO SAPIENS |
| 1Y64 | Actin | BNI1 protein | OX | 2745 | SACCHAROMYCES CEREVISIAE; ORYCTOLAGUS CUNICULUS |
| 1YVB | Falcipain 2 | Cystatin | EI | 1743 | PLASMODIUM FALCIPARUM; GALLUS GALLUS |
| 1Z0K | Rab4A GTPase | RAB4 binding domain of Rabenosyn | OG | 1787 | HOMO SAPIENS |
| 1Z5Y | N-term of DsbD | E.coli CCMG protein | ES | 1346 | ESCHERICHIA COLI |
| 1ZHH | Autoinducer 2-binding periplasmic protein LuxP | Autoinducer 2 sensor kinase/phosphatase LuxQ | OR | 2189 | VIBRIO HARVEYI |
| 1ZHI | BAH domain of Orc1 | Sir Orc-interaction domain | OX | 1322 | SACCHAROMYCES CEREVISIAE |
| 1ZLI | Carboxypeptidase B | Tick carboxypeptidase inhibitor | EI | 2084 | HOMO SAPIENS; RHIPICEPHALUS BURSA |
| 1ZM4 | Elongation factor 2 | Diphtheria toxin A catalytic domain | ES | 1554 | PSEUDOMONAS AERUGINOSA; SACCHAROMYCES CEREVISIAE |
| 2A1A | Eukayotic translation initiation factor 2-alpha kinase 2 | eIF2 alpha subunit | ES | 1186 | SACCHAROMYCES CEREVISIAE; HOMO SAPIENS |
| 2A5T | NMDA receptor R1-4A subunit ligand-binding core | NMDA receptor R2A subunit ligand-binding core | OX | 1892 | RATTUS NORVEGICUS; CANIS LUPUS FAMILIARIS |
| 2A9K | Ras-related protein Ral-A | Mono-ADP-ribosyltransferase C3 | ES | 1751 | HOMO SAPIENS; CLOSTRIDIUM BOTULINUM D PHAGE |
| 2ABZ | Carboxypeptidase A1 | Leech carboxypeptidase inhibitor | EI | 1443 | BOS TAURUS; HIRUDO MEDICINALIS |
| 2AJF | ACE2 | SARS spike protein receptor binding domain | OR | 1704 | HOMO SAPIENS; SARS CORONAVIRUS |
| 2AYO | Ubiquitin carboxyl-terminal hydrolase 14 | Ubiquitin | ER | 3027 | HOMO SAPIENS |
| 2B42 | Xylanase | Xylanase inhibitor | EI | 2520 | BACILLUS SUBTILIS; TRITICUM AESTIVUM |
| 2BTF | Actin | Profilin | OX | 2063 | BOS TAURUS |
| 2C0L | PTS1 and TRP region of PEX5 | SCP2 | OX | 2013 | HOMO SAPIENS |
| 2CFH | BET3 | TPC6 | OX | 2384 | HOMO SAPIENS |
| 2FJU | Phospholipase Beta 2 | Rac GTPase | OG | 1245 | HOMO SAPIENS |
| 2G77 | GTPase-activating protein GYP1 | Ras-related protein Rab-33B | OG | 2524 | SACCHAROMYCES CEREVISIAE; MUS MUSCULUS |
| 2GAF | Poly(A) polymerase VP55 | Vaccinia protein VP39 | ER | 3368 | VACCINIA VIRUS |

| | | | | | |
|---|---|---|---|---|---|
| 2GTP | Alpha-1 subunit Guanine nucleotide-binding protein G(I), alpha-1 subunit | RGS1 | OG | 1442 | HOMO SAPIENS |
| 2H7V | Rac GTPase | YpkA | OG | 1574 | HOMO SAPIENS; YERSINIA PSEUDOTUBERCULOSIS |
| 2HLE | Ephrin B4 receptor | Ephrin B2 ectodomain | OR | 2116 | HOMO SAPIENS |
| 2HQS | TolB | Pal | OX | 2333 | ESCHERICHIA COLI |
| 2HRK | Glutamyl-t-RNA synthetase | GU-4 nucleic binding protein | OX | 1595 | SACCHAROMYCES CEREVISIAE |
| 2I25 | Shark single domain antigen receptor | Lysozyme | AA | 1425 | GINGLYMOSTOMA CIRRATUM; GALLUS GALLUS |
| 2I9B | Urokinase plasminogen activator surface receptor | Urokinase-type plasminogen activator | OR | 2371 | HOMO SAPIENS |
| 2IDO | DNA polymerase III Œ$\mu$ exonuclease domain | HOT protein (P1 phage) | ES | 1953 | ESCHERICHIA COLI; ENTEROBACTERIA PHAGE |
| 2J0T | MMP1 Intersitial collagenase | Metalloproteinase inhibitor 1 | EI | 1477 | HOMO SAPIENS |
| 2J7P | SRP GTPase Ffh | Cell division protein FtsY | OX | 3008 | THERMUS AQUATICUS |
| 2NZ8 | Rac GTPase | DH/PH domain of TRIO | ER | 2599 | HOMO SAPIENS |
| 2O3B | NucA nuclease | NuiA nuclease inhibitor | EI | 1675 | NOSTOC SP. |
| 2O8V | PAPS reductase | Thioredoxin | ES | 1619 | ESCHERICHIA COLI |
| 2OOB | Ubiquitin ligase | Ubiquitin | ES | 808 | HOMO SAPIENS; BOS TAURUS |
| 2OT3 | Rab21 GTPase | Rabex-5 VPS9 domain | ER | 2306 | HOMO SAPIENS |
| 2OUL | Falcipain 2 | Chagasin | EI | 1933 | PLASMODIUM FALCIPARUM; TRYPANOSOMA CRUZI |
| 2OZA | MAP kinase 14 | MAP kinase-activated protein kinase 2 | OX | 6248 | HOMO SAPIENS; MUS MUSCULUS |
| 2PCC | Cyt C peroxidase | Cytochrome C | ES | 1141 | SACCHAROMYCES CEREVISIAE |
| 2SIC | Subtilisin | Streptomyces subtilisin inhibitor | EI | 1617 | BACILLUS AMYLOLIQUEFACIENS |
| 2SNI | Subtilisin | Chymotrypsin inhibitor 2 | EI | 1628 | BACILLUS AMYLOLIQUEFACIENS; HORDEUM SP. |
| 2UUY | Trypsin | Tryptase inhibitor from tick | EI | 1280 | BOS TAURUS; RHIPICEPHALUS APPENDICULATUS |
| 2VDB | Serum albumin | Peptostreptococcal albumin-binding protein | OX | 1798 | PEPTOSTREPTOCOCCUS MAGNUS; HOMO SAPIENS |
| 2X9A | TolA C-terminal domain | G3P TolA binding domain | OR | 1571 | ENTEROBACTERIA PHAGE IF1; ESCHERICHIA COLI |
| 2YVJ | Ferredoxin reductase BPHA4 | Biphenyl dioxygenase ferredoxin subunit | ER | 1377 | PSEUDOMONAS SP. |
| 2Z0E | Cysteine protease Atg4B | Microtubule-associated proteins 1A/1B light chain 3B | ER | 2478 | HOMO SAPIENS; RATTUS NORVEGICUS |
| 3A4S | SUMO-conjugating enzyme UBC9 | NFATC2-interacting protein SLD2 ubiquitin-like domain | EI | 1116 | HOMO SAPIENS; MUS MUSCULUS |
| 3AAD | Double bromodomain | Histone chaperone ASF1 | OX | 1654 | HOMO SAPIENS |

| | | | | | |
|---|---|---|---|---|---|
| 3BIW | Neuroligin-1 | Neuroligin-1-beta | OX | 1191 | RATTUS NORVEGICUS |
| 3BX7 | Lipocalin 2 | CTLA-4 extracellular domain | OX | 2349 | HOMO SAPIENS |
| 3CPH | Rab GDP-dissociation inhibitor | Ras-related protein Sec4 | OG | 1685 | SACCHAROMYCES CEREVISIAE |
| 3D5S | Complement C3d fragment | Fibrinogen-binding protein C-ter domain | OX | 1620 | HOMO SAPIENS; STAPHYLOCOCCUS AUREUS SUBSP. AUREUS STR. NEWMAN |
| 3DAW | Alpha actin | Twinfilin-1 C-terminal domain | OX | 2323 | MUS MUSCULUS; ORYCTOLAGUS CUNICULUS |
| 3F1P | HIF2 alpha C-terminal PAS domain | ARNT C-terminal PAS domain | OX | 1919 | HOMO SAPIENS |
| 3FN1 | UQ_con domain from NEDD8-conjugating enzyme UBE2F | NEDD8-activaating enzyme E1 catalytic subunit | ER | 1897 | HOMO SAPIENS |
| 3H2V | Vinculin tail domain | Raver1 RRM1 domain | OX | 1263 | HOMO SAPIENS |
| 3K75 | DNA polymerase beta | Reduced XRCC1, N-terminal domain | ER | 1195 | HOMO SAPIENS; RATTUS NORVEGICUS |
| 3PC8 | DNA repair protein XRCC1 | DNA ligase III-alpha BRCT domain | ER | 1240 | MUS MUSCULUS; HOMO SAPIENS |
| 3RJQ | A12 | C1086 HIV gp120 | AA | 1734 | HUMAN IMMUNODEFICIENCY VIRUS TYPE 1; LAMA GLAMA |
| 3S9D | IFNAR2 | IFNa2 | OR | 1841 | HOMO SAPIENS |
| 3SGQ | Streptogrisin B | Ovomucoid inhibitor third domain | EI | 1211 | MELEAGRIS GALLOPAVO; STREPTOMYCES GRISEUS |
| 3VLB | EDGP | Xyloglucan-specific endo-beta-1,4-glucanase A | EI | 2020 | DAUCUS CAROTA; ASPERGILLUS ACULEATUS |
| 4CPA | Carboxypeptidase A | Potato carboxypeptidase inhibitor | EI | 1175 | BOS TAURUS; SOLANUM TUBEROSUM |
| 4FZA | MO25 alpha | Serine/threonine-protein kinase MST4 | ER | 1695 | HOMO SAPIENS |
| 4H03 | Iota toxin component IA | Alpha actin | ES | 1474 | CLOSTRIDIUM PERFRINGENS; ORYCTOLAGUS CUNICULUS |
| 4IZ7 | Non-phosphorylated ERK | PEA-15 Death Effector Domain | EI | 1202 | HOMO SAPIENS; CRICETULUS GRISEUS |
| 4M3K | cAb-H7S | B. licheniformis beta-lactamase | AA | 1588 | BACILLUS LICHENIFORMIS; LAMA GLAMA |
| 4M76 | C3D | Integrin alpha-M CD11B A-domain | OR | 1046 | HOMO SAPIENS |
| 4POU | VHHmetal | bovine RNase A | AA | 1313 | LAMA GLAMA; BOS TAURUS |
| 4Y7M | nb25 | E coli TssM CTD | AA | 1103 | LAMA GLAMA; ESCHERICHIA COLI 2-156-04_S3_C3 |
| 5E5M | H11 | mouse CTLA-4 | AA | 1341 | MUS MUSCULUS; CAMELIDAE |
| 5HGG | Nb4 | uPA | AA | 1969 | HOMO SAPIENS; VICUGNA PACOS |
| 5JMO | Nb14 | Furin | AA | 1394 | HOMO SAPIENS; CAMELUS DROMEDARIUS; SYNTHETIC CONSTRUCT |
| 5SV3 | A3C8 | Ricin | AA | 1294 | LAMA GLAMA; RICINUS COMMUNIS |
| 5VNW | Nb.b201 | human serum albumin | AA | 967 | HOMO SAPIENS; SYNTHETIC CONSTRUCT |
| 6CWG | A9 | Ricin | AA | 1151 | RICINUS COMMUNIS; VICUGNA PACOS |

| | | | | | |
|---|---|---|---|---|---|
| 6DBG | R303 | Listeria monocytogenes internalin B | AA | 1525 | LISTERIA MONOCYTOGENES; CAMELUS DROMEDARIUS |
| 7CEI | Colicin E7 nuclease | Im7 immunity protein | EI | 1384 | ESCHERICHIA COLI STR. K12 SUBSTR. |
| BAAD | Double bromodomain | Histone chaperone ASF1 | OX | 1461 | HOMO SAPIENS |
| BOYV | Subtilisin Carlsberg | Two-headed tomato inhibitor-II | EI | 1280 | BACILLUS LICHENIFORMIS; SOLANUM LYCOPERSICUM |

[a] Complex category: Antibody-antigen (AA), enzyme-inhibitor (EI), enzyme-substrate (ES), enzyme complex with a regulatory or accessory chain (ER), others, G-protein containing (OG), others, receptor-containing (OR); others, miscellaneous (OX).
[b] Interface buried surface area was obtained from the BM5.5 site (https://zlab.umassmed.edu/benchmark/).
[c] Source organisms for receptor and ligand protein chains in the Protein Data Bank (PDB) [3], separated by semicolon. If both the ligand and receptor proteins in the complex were from the same source organism, only one source organism is shown.

**Table S2.** Area under the ROC curve (AUC) value for protein quality classes as a function of interface scores calculated from AlphaFold predictions.

| Score[a] | Binary classification | | Multi-class classification |
|---|---|---|---|
| | Incorrect vs. High | Incorrect vs. Medium and High | |
| Interface PAE (4 Å) | 0.93 | 0.90 | 0.81 |
| Interface PAE (5 Å) | 0.92 | 0.89 | 0.81 |
| Interface PAE (6 Å) | 0.91 | 0.89 | 0.80 |
| Interface PAE (7 Å) | 0.91 | 0.88 | 0.80 |
| Interface PAE (10 Å) | 0.91 | 0.88 | 0.80 |
| Interface pLDDT (4 Å) | 0.97 | 0.90 | 0.84 |
| Interface pLDDT (5 Å) | 0.95 | 0.88 | 0.82 |
| Interface pLDDT (6 Å) | 0.94 | 0.85 | 0.80 |
| Interface pLDDT (7 Å) | 0.93 | 0.83 | 0.77 |
| Interface pLDDT (10 Å) | 0.91 | 0.81 | 0.77 |

[a] Scoring methods. "Interface PAE": the average PAE score of pairs of interface residues within the interface distance cutoff specified in parenthesis; "Interface pLDDT": the average pLDDT of interface residues within the interface distance cutoff specified in parenthesis.

**Table S3. Additional antibody-antigen complex test cases and AlphaFold prediction success.**

| PDB | Antibody | Antigen | Type[a] | AlphaFold T1[b] | AlphaFold T5[b] |
|---|---|---|---|---|---|
| 5F72 | LS146 | KEAP1 | ab | Incorrect | Incorrect |
| 1DZB | 1F9 | HEW | ab | Incorrect | Incorrect |
| 3UZE | 4E11 | Dengue E DIII | ab | Incorrect | Incorrect |
| 6EJM | scFv 5 | CD81 LEL | ab | Incorrect | Incorrect |
| 5DFW | K13 | CD81 LEL | ab | Incorrect | Incorrect |
| 6I07 | MM131 | EpCAM | ab | Incorrect | Incorrect |
| 5JYM | TSP11 | P-cadherin | ab | Incorrect | Incorrect |
| 4NIK | F5 | Gankyrin | ab | Medium | Medium |
| 5JYL | TSP7 | P-cadherin | ab | Incorrect | Incorrect |
| 6TOU | RVC20 | Rabies gp | ab | Incorrect | Incorrect |
| 6EK2 | scFv 10 | CD81 LEL | ab | Incorrect | Incorrect |
| 4YJZ | H2526 | H1 HA | ab | Incorrect | Incorrect |
| 3UX9 | AIFNα1bScFv01 | interferon alpha | ab | Incorrect | Incorrect |
| 6OAN | Antibody 053054 | P vivax DBP | ab | Medium | Medium |
| 6J71 | HUA21 | HER2 | ab | Incorrect | Incorrect |
| 7DET | PR961 | SARS-CoV-2 RBD | ab | Incorrect | Incorrect |
| 7DEO | PR1077 | SARS-CoV-2 RBD | ab | Incorrect | Incorrect |
| 6WAQ | VHH-72 | SARS-CoV RBD | nano | Incorrect | Incorrect |
| 6EY0 | NB01 | PorM | nano | Incorrect | Incorrect |
| 6OQ7 | E3 | Clostridium difficile toxin B | nano | Incorrect | Incorrect |

[a] Antibody type: "ab": heavy-light chain antibody, "nano": nanobody/VHH.
[b] AlphaFold modeling success in top 1 and top 5 models (ranked by pTM). Model quality is assessed by CAPRI criteria.
